# Molecular Basis of Kv1 Channel Downregulation and Its Epileptogenic Implications in Lgi1 Knock-out Mice

**DOI:** 10.1101/2023.07.24.550266

**Authors:** Jorge Ramirez-Franco, Kévin Debreux, Marion Sangiardi, Maya Belghazi, Yujin Kim, Suk-Ho Lee, Christian Lévêque, Michael Seagar, Oussama El Far

## Abstract

The Kv_1_ members (KCNA, Shaker) of the voltage-gated potassium channels are implicated in determining key functional neuronal properties from spike generation at axonal initial segments to the control of synaptic strength at nerve terminals. In animal models of LGI1-dependent autosomal dominant lateral temporal lobe epilepsy (ADTLE), Kv_1_ channels are downregulated, suggesting their crucial involvement in epileptogenesis. The molecular basis of Kv_1_ channel-downregulation in LGI1 knock-out mice has not been elucidated and how the absence of this extracellular protein induces an important modification in the expression of Kv_1_ remains unknown. In this study we analyse by immunofluorescence the detailed modifications in neuronal Kv_1.1_ and Kv_1.2_ distribution throughout the hippocampal formation of LGI1 knock-out mice. We show that Kv_1_ downregulation is not restricted to the axonal compartment, but also takes place in the somatodendritic region and is accompanied by a drastic decrease in Kv_2_ expression levels. Moreover, we find that the downregulation of these Kv channels is associated with an important increase in bursting patterns. Finally, mass spectrometry uncovered key important modifications in the Kv_1_ interactome that highlight the epileptogenic implication of Kv_1_ downregulation in LGI1 knock-out animals.

## 1. Introduction

The delayed rectifying Kv_1_ channels which mediate low voltage-activated K^+^ currents are key regulators of neuronal excitability, affecting action-potential (AP) threshold and firing patterns.^[1 2 3]^ Functional plasticity in neurons involves dynamic changes in the expression and subcellular distribution of potassium channels at distinct neuronal compartments.^[4]^ Kv_1_ channels are formed by homomeric or heteromeric co-assembly of four alpha subunits with auxiliary subunits.^[4]^ A subset of Kv_1_ subunits, namely Kv_1.1_ (KCNA1), Kv_1.2_ (KCNA2) and Kv_1.6_ (KCNA6) are responsible for the α-dendrotoxin (α-DTX) sensitive D-type current and these Kv subunits are present in axonal and synaptic compartments ^[5 6]^ as well as at axonal initial segments.^[7 8]^

Gain-or loss-of-function, inherited or de novo missense variants of these subunits lead to neurological disorders which display extensive clinical heterogeneity.^[9 10 11 12]^ LGI1 is a secreted extracellular glycoprotein^13^ which is associated with Kv_1_ channels.^[14 15 16]^ Mutations in the LGI1 gene often result in the inhibition of its secretion^[13 17]^which underlies hyperexcitability in Autosomal Dominant Lateral Temporal Lobe Epilepsy (ADLTE). Also, the production of LGI1-autoantibodies promotes neuronal excitability and epilepsy in LGI1-dependent autoimmune limbic encephalitis. In both genetic and autoimmune disruption of LGI1 function, a significant decrease in total, as well as plasma membrane, Kv_1.1_ and Kv_1.2_ is observed.^[8 18 19]^ This decrease may underlie the increase in neuronal excitability in ADLTE^[8]^ and precedes the previously described ^[20]^ perturbation in AMPA/NMDA receptors ratio.^[21 8]^ An additional secretion-defective LGI1 mutation was recently found to induce Kv_1.1_ downregulation underlying neuronal hyperexcitability and irregular spiking in the CA1 pyramidal cells.^[22]^ Although LGI1 is part of the Kv_1_-associated proteome ^[14 15 16 23]^, how the absence of an extracellular glycoprotein leads to the downregulation of Kv_1_ subunits has not yet been elucidated. LGI1 binds to the proteolytically-inactive transmembrane ADAM family members ADAM11, 22 and 23 ^[24 25 26]^ and Kv_1_ is a constituent of the ADAM22-associated proteome ^[20]^. While ADAM22^[17]^ as well as Kv_1.1_ and Kv_1.2_ C-terminal tails, directly interact with PSD95/93 via PDZ domains^[27]^, the interplay between ADAM, PSD95/93 and Kv1 subunits is complex. PSD93 (DLG2), but not PSD95 (DLG4) ^[28 14]^ nor ADAM22 ^[14]^, is important for Kv_1.1_ and Kv_1.2_ channel localisation and clustering at axonal initial segments.^[5]^ Also, PDZ interaction domains of Kv_1_ are crucial for surface expression but not axonal targeting.^[6]^

In this study we unveil the molecular mechanisms by which the absence of LGI1 triggers a decrease in the expression and mislocalization of several potassium channels at different cellular loci throughout the hippocampal formation. The use of mass spectrometry uncovered key modifications in the Kv_1_ interactome that highlight the epileptogenic implication of Kv1 downregulation. We finally show that complex bursting patterns dominate in *Lgi1^-/-^* CA3 pyramidal neurons, and that this is associated with a substantial decrease in the somatodendritic Kv_2_ channels that are known to participate in modulating spike bursting patterns.^[29]^

## 2. Results

### 2.1. Kv_1.1_ and Kv_1.2_ expression in *Lgi1^-/-^*

We have previously shown that Kv_1.1_ and Kv_1.2_ subunits were massively downregulated in LGI1 knock-out mice (*Lgi1^-/-^*) total brain extracts and that, in this genetic background, Kv_1.1_ was almost absent from axonal initial segments of hippocampal and cortical neurons^[8]^. In the light of the major enrichment of LGI1 in the hippocampal formation ^[23]^, we compared by Western blot the downregulation level of Kv_1.1_ and Kv_1.2_ subunits in total brain homogenate as well as in hippocampal extracts **(Figure 1 and 2)**. We observed that the decrease in expression of Kv_1.1_ and Kv_1.2_ in total brain (Figure 1a) (Kv_1.1_ WT 1.00 ± 0.07 vs Kv_1.1_ *Lgi1^-/-^* 0.72 ± 0.08, p = 0.047; Kv_1.2_ WT 1.00 ± 0.05 vs Kv_1.2_ *Lgi1^-/-^*0.52 ± 0.04, p = 1.03 x 10^-7^) was less severe than in hippocampus (Figure 1b, Kv_1.1_ WT 1.00 ± 0.05 vs Kv_1.1_ *Lgi1^-/-^* 0.55 ± 0.03, p =1.33 x 10^-7^; Kv_1.2_ WT 1.00 ± 0.04 vs Kv_1.2_ *Lgi1^-/-^* 0.38 ± 0.02, p = 1.73 x 10^-13^).

**Figure 1.**
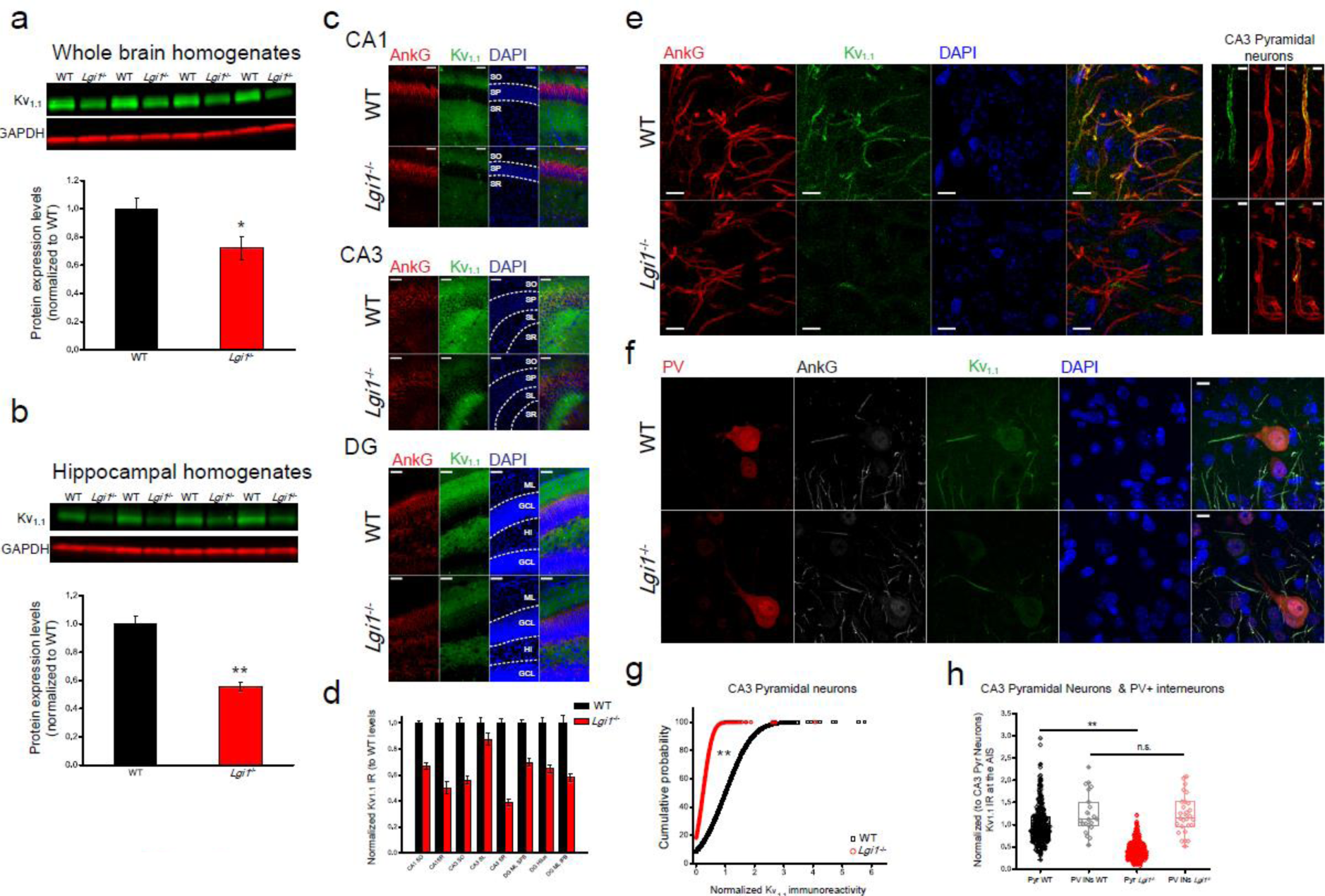
Kv_1.1_ expression is reduced in *Lgi1*^-/-^ mice. A) Quantification by Western blot of Kv_1.1_ expression in whole brain homogenates of WT (1.00 ± 0.07) and *Lgi1*^-/-^ (0.72 ± 0.08), p = 0.047; n = 4; Two-sample t-test). b) Quantification by Western blot of Kv_1.1_ expression in hippocampal homogenates of WT (1.00 ± 0.05) and *Lgi1^-/-^* (0.55 ± 0.03); p = 1.33 x 10^-7^; n =1 6; Two-sample t-test). In a and b, signals were normalized to GAPDH expression levels. c) Representative images of Kv_1.1_ immunoreactivity (IR) across CA1 (top panels, Scale bar = 50 µm), CA3 (middle panels, Scale bar = 50 µm), and DG (bottom panels, Scale bar = 50 µm) in WT vs *Lgi1*^-/-^ mouse. SO: Str. Oriens; SP: Str. Pyramidale; SR: Str. Radiatum; SL: Str. Lucidum; ML: Mol. Layer; GCL: Granule Cell Layer; Hi: Hilus. d) Quantification of the differences in the normalized Kv_1.1_ IR (to WT levels) through the different hippocampal strata. For the sake of clarity, only non-significant comparisons are indicated. For all other comparison \*\**p* < 0.01 (CA1-SO WT = 1.00 ± 0.02 vs *Lgi1*^-/-^ = 0.66 ± 0.02 p = 5.96×10^-13^; CA1-SR WT = 1.00 ± 0.03 vs *Lgi1*^-/-^ = 0.05 ± 0.05 p = 1.20×10^-6^; CA3-SO WT = 1.00 ± 0.04 vs *Lgi1*^-/-^ = 0.53 ± 0.03 p = 2.75×10^-10^; CA3-SL WT =1.00 ± 0.05 vs *Lgi1*^-/-^ = 0.88 ± 0.05 p = 0.07; CA3-SR WT = 1.00 ± 0.03 vs *Lgi1*^-/-^= 0.39 ± 0.02 p = 2.34×10^-10^; DG-ML-SPB WT=1.00 ± 0.02 vs *Lgi1*^-/-^ = 0.70 ± 0.03 p = 1.20 x 10^-8^; DG-Hilus WT = 1.00 ± 0.04 vs *Lgi1*^-/-^ = 0.65 ± 0.03 p = 3.57×10^-8^; DG-ML-IPB WT = 1.00 ± 0.06 vs *Lgi1*^-/-^ = 0.58 ± 0.03 p = 1.03×10^-7^; Two-sample t-test). e) Left, 63x images of Kv_1.1_ immunoreactivity at CA3 AIS in WT (top panels) and *Lgi1^-/-^*(bottom panels). Scale bar = 10 µm. Right, detail of a CA3 pyramidal neuron AIS in WT (top panels) and *Lgi1^-/-^* (bottom panels). Scale bar = 2.5 µm. f) 63x images showing no differences in Kv_1.1_ IR at the AIS of PV+ interneurons g) Cumulative probability plots of Kv_1.1_ immunoreactivity at individual AIS of WT (black squares) and *Lgi1*^-/-^ (red circles) CA3 pyramidal neurons (WT Pyr CA3 neurons= 1.00 ± 0.02 vs *Lgi1*^-/-^ CA3 Pyr neurons = 0.23 ± 0.01; p = 0.00, Kolmogorov-Smirnov Test). h) Comparisons of Kv_1.1_ immunoreactivity (Normalized to WT CA3 Pyr neurons) at the AIS of pyramidal cells (dark tones) and PV+ interneurons (light tones) across WT (black) and *Lgi1*^-/-^ (red) hippocampi (Pyr WT = 1.00 ± 0.02 vs Pyr *Lgi1*^-/-^ = 0.40 ± 0.01 p = 2.43×10^-132^; PV+ IN WT = 1.25 ± 0.10 vs PV+ IN *Lgi1*^-/-^ = 1.22 ± 0.08; p = 1.00; ANOVA followed by Bonferroni’s test for means comparison).

**Figure 2.**
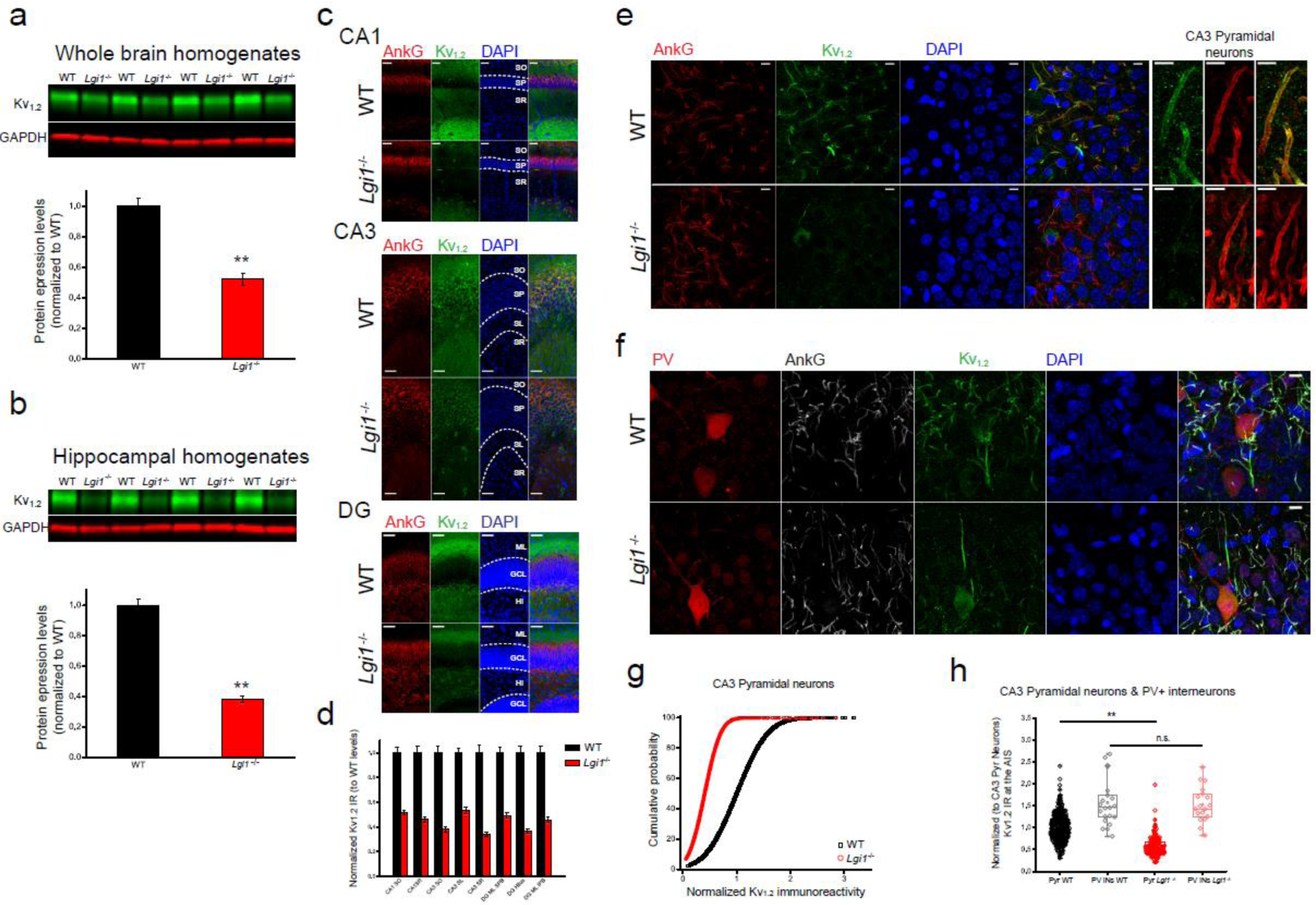
Kv_1.2_ expression is reduced in the *Lgi1*^-/-^ mice. a) Quantification by Western blot of Kv1.2 expression in brain homogenates of WT (1.00 ± 0.06) and *Lgi1*^-/-^ (0.52 ± 0.04), p = 1.03 x 10^-7^; n = 17; Two-sample t-test). b) Quantification by Western blot of Kv_1.2_ in in hippocampal homogenates of WT (1.00 ± 0.04) and *Lgi1^-/-^* (0.38 ± 0.02); p = 1.73 x 10^-13^; n = 20; Two-sample t-test). In **a** and **b**, signals were normalized to GAPDH expression levels. c) representative images of Kv_1.2_ immunoreactivity (IR) across CA1 (top panels, Scale bar = 50 µm), CA3 (middle panels, Scale bar = 50 µm), and DG (bottom panels, Scale bar = 50 µm) in WT vs *Lgi1*^-/-^ mouse. SO: Str. Oriens; SP: Str. Pyramidale; SR: Str. Radiatum; SL: Str. Lucidum; ML: Mol. Layer; GCL: Granule Cell Layer; Hi: Hilus. d) Quantification of the differences in the normalized Kv_1.2_ IR (to WT levels) through the different hippocampal strata, all comparisons are \*\**p* < 0.01 (CA1-SO WT = 1.00 ± 0.04 vs *Lgi1*^-/-^ = 0.52 ± 0.02 p = 1.58×10^-13^; CA1-SR WT = 1.00 ± 0.06 vs *Lgi1*^-/-^ = 0.46 ± 0.02 p = 8.37×10^-10^; CA3-SO WT = 1.00 ± 0.06 vs *Lgi1*^-/-^ = 0.38 ± 0.02 p = 6.32×10^-11^; CA3-SL WT = 1.00 ± 0.04 vs *Lgi1*^-/-^ = 0.53 ± 0.03 p = 9.61×10^-13^; CA3-SR WT = 1.00 ± 0.06 vs *Lgi1*^-/-^ = 0.34 ± 0.02 p = 7.49 x 10^-11^; DG-ML-SPB WT = 1.00 ± 0.05 vs *Lgi1*^-/-^ = 0.49 ± 0.02 p = 1.92 x 10^-11^; DG-Hilus WT = 1.00 ± 0.05 vs *Lgi1*^-/-^ = 0.37 ± 0.01 p = 4.15 x 10^-12^; DG-ML-IPB WT = 1.00 ± 0.05 vs *Lgi1*^-/-^ = 0.46 ± 0.02 p = 1.83 x 10^-10^; Two-sample t-test). e) Left, 63x images of Kv1.2 immunoreactivity at CA3 Axonal Initial segments in WT (top panels) and *Lgi1*^-/-^ (bottom panels). Scale bar = 10 µm. Right, detail of a CA3 pyramidal neuron AIS in WT (top panels) and *Lgi1^-/-^* (bottom panels). Scale bar = 2.5 µm. f) 63x images showing no differences in Kv_1.2_ IR at the AIS of PV+ interneurons. g) Cumulative probability plots of Kv_1.2_ immunoreactivity at individual AIS of WT (black squares) and *Lgi1*^-/-^ (red circles) CA3 pyramidal neurons (WT Pyr CA3 neurons= 1.00 ± 0.01 vs *Lgi1*^-/-^ CA3 Pyr neurons = 0.41 ± 0.01; p = 0.00, Kolmogorov-Smirnov Test). h) Comparisons of Kv_1.2_ immunoreactivity (Normalized to WT CA3 Pyr Neurons) at the AIS of pyramidal cells (dark tones) and PV+ interneurons (light tones) across WT (black) and *Lgi1*^-/-^ (red) hippocampi (Pyr WT = 1.00 ± 0.02 vs Pyr *Lgi1*^-/-^ = 0.55±0.01 p = 7.90×10^-60^; PV+ IN WT=1.57 ± 0.12 vs PV+ IN *Lgi1*^-/-^ =1.53 ± 0.09; p = 1.00; ANOVA followed by Bonferroni’s test for means comparison).

In order to address Kv_1_ expression and distribution profile, we investigated, the detailed comparative immunofluorescence staining of Kv_1.1_ and Kv_1.2_ in the hippocampal formation in both WT and *Lgi1^-/-^* backgrounds. It is noteworthy that conventional immunohistochemical techniques could render the epitopes of Kv_1.1_ and Kv_1.2_ inaccessible to antibodies, especially in slices from prefixed brain, precluding the detection of these channels particularly in the AIS of hippocampal neurons ^[30]^. In our hands, this proved to be the case, since Kv_1.1_ was not detected in the absence of antigen retrieval protocols (Figure S1, Supporting Information). The use of the same antibody on organotypic slices ^[31]^ or using mild fixation conditions in thin slices ^[21]^ could facilitate epitope access. Due to the tortuosity of the AIS in CA3, we chose to use 50 μm thick sections and therefore, employing antigen retrieval techniques ^[32 33]^ and addressed Kv_1.1_ (Figure 1c-h) and Kv_1.2_ (Figure 2c-h) distribution. In addition to the previously reported decrease in Kv_1_ expression in the AIS (Figure 1e and Figure 2e), we observed an overall decrease in all hippocampal strata (CA1, CA3 and DG) (Figure 1c,d,g and Figure 2c,d,g) with a more prominent decrease of Kv_1.2_ versus Kv_1.1_. Interestingly, we observed that in parvalbumin (PV) positive interneurons, the expression levels of both Kv_1.1_ and Kv_1.2_ subunits were not affected, (Figure 1f,h and Figure 2f,h) suggesting that Kv_1_ expression levels in PV+ cells are not regulated by LGI1.

### 2.2. Characterization of Kv1.2 antibodies

In order to address changes in Kv_1.2_ proteome between WT and *Lgi1^-/-^*, we generated a polyclonal antibody directed against a C-terminal peptide sequence and performed immunoprecipitations followed by mass spectrometry analysis. As shown in Figure S2a Supporting Information, this antibody recognizes Kv_1.2_ from brain extracts as well as recombinant pHLuorin-tagged full length Kv_1.2_ but not Kv_1.1_ expressed in HEK cells. However, the band recognized in wild-type mice brain extracts was still present in Kv_1.2_ knock out extracts (Figure S2b Supporting Information) suggesting that this antibody can cross-react, in native tissues, with closely homologous Kv_1_ isoforms. Due to the significant sequence homology between the target peptide sequence in other Kv_1_ family members (Figure S3a, Supporting Information), we verified whether at higher protein concentrations, the antibody can recognize other Kv_1_ family members. As shown in Figure S3b,c Supporting Information, the antibody showed cross-reactivity with GST fusion proteins containing C-terminal sequences of Kv_1.1_ and to a lesser extent Kv_1.3_ but not Kv_1.4_. These results indicate that depending on the abundance of the Kv_1_ isoform, the used antibody can cross reacts with other closely related and significantly homologous Kv_1_ family members.

### 2.3 Kv_1_-associated proteome in WT versus *Lgi1*^-/-^ brain homogenate

The capacity of our antibody to recognize Kv_1.1_ and Kv_1.2_ allows us to address the consequences of the absence of LGI1 on the modification of the proteome associated with these hetero-multimeric potassium channels. Kv_1_α (KCNA1 and 2) subunits as well as other Kvβ1 (KCAB1) and β2 (KCAB2) were equally recovered except Kv_1.6_ (KCNA6) that was less immunoprecipitated (−32,8%) from *Lgi1^-/-^ extracts* (Table S1, and Supplementary sheet). The comparison between Kv_1_ partners in WT and *Lgi1^-/-^* reveals intriguing findings, suggesting a dynamic interplay of protein interactions. Certain partners are lost in the absence of LGI1, while new partners are acquired, highlighting the remodelling of Kv_1_ interaction networks. Furthermore, the analysis indicates that the interaction with specific partners is enhanced, indicating a potential compensatory mechanism, while the association with other partners is diminished.

#### 2.3.1. Lost Kv_1_ partners in Lgi1^-/-^ samples

The most striking observation was that the landscape of the Kv_1_ links with the cytoskeleton shows major rearrangements (Supplementary Tables 2, 3, Figure S4 Supplementary Information and Supplementary sheet). The association with ACTC Actin (alpha cardiac muscle 1), the most abundant Kv_1_ partner from WT extracts is lost in the *Lgi1^-/-^* samples. This actin isoform was shown to be widely expressed in the hippocampus and other brain structures. ^[34]^ In addition to actin, the co-immunoprecipitation of myosin 10 and the intermediate filament vimentin as well as the actin remodelling protein CYFP2 (cytoplasmic FMR1 Interacting Protein 2 also known as PIR12) ^[35 36]^ are completely disrupted in *Lgi1^-/-^*. Interestingly, CYFP2 is linked to mTORC signalling ^[37]^ and was shown to be involved in epileptic encephalopathy and intellectual disability.^[38]^ The perturbations in membrane expression and cytoskeletal links of Kv_1_ are also reflected by the loss of interaction of Kv_1_ with the following partners: i-the membrane-associated phosphatidylinositol (4,5)-bisphosphate binding protein GSDMA, that has pore forming properties and is involved in cell death ^[39]^, ii-the deltex E3 ubiquitin-protein ligase and regulator of Notch signalling DTX1, iii-the multi-modular scaffolding protein RANB9 that is implicated in actin cytoskeletal rearrangement ^[40]^ and the integration of a variety of cell surface receptor signalling through intracellular targets ^[41]^, iv-The PM/ER cholesterol transporter and phosphatidylserine binding protein ^[42]^ ASTRA that may indicate a change in the lipidic environment of Kv_1_. Although a similar function was attributed to ASTRA and B, the decrease of interaction with Kv_1_ is restricted to ASTRA, ASTRB remaining stable, v-the NEB2 (spinophilin) ^[43]^, a PDZ-domain containing F-actin binding and phosphatase recruiter scaffolding protein. NEB2 is a regulatory subunit of protein phosphatase 1 ^[43]^ that controls serine/threonine dephosphorylation of several membrane proteins such as ion channels and certain G-protein coupled receptors ^[44]^ and is enriched in postsynaptic compartments ^[45]^ where it is involved in modulating spine maturation and morphology.^[46]^

In accordance with this latter observation, the association of Kv_1_ with PP2BB is also lost in *Lgi1^-/-^*samples. PP2BB is a regulatory subunit of the activity-dependent serine/threonine-protein phosphatase calcineurin and plays an essential role in intracellular Ca^2+^-mediated signal transduction. With PP1, it plays a regulatory role in Kv_2.1_ (see later for the description of Kv_2_ decrease) phosphorylation modulating its activity, membrane clustering and internalisation with key effects on the control of neuronal excitability ^[47 44]^. Interestingly, the interaction of NEB2 with actin is modulated by doublecortin (DCX) ^[48]^ and we found that this protein is acquired as a new partner of Kv_1_ in *Lgi1^-/-^* samples.

Intriguingly, a specific loss of association with PSD93 but not with its twin PSD95 (DLG4) or other MAGUKs is observed in *Lgi1^-/-^*. We therefore investigated by Western blot and immunofluorescence whether the loss of association with PSD93 is accompanied by a decrease in its total expression level. As shown in Figure 3, PSD93 expression levels were severely downregulated in *Lgi1^-/-^* samples with a more prominent decrease in total brain homogenates (PSD93 WT 1.00 ± 0.03 vs PSD93 *Lgi1*^-/-^ 0.46 ± 0.04, p = 1.40×10^-11^) (Figure 3a) compared to hippocampal extracts (PSD93 WT 1.00 ± 0.05 vs PSD93 *Lgi1*^-/-^ 0.68 ± 0.05, p = 6.10×10^-5^) (Figure 3b). A closer look at the subcellular distribution of PSD93 indicates that this decrease was not restricted to the AIS (Figure 3e,f) and was observed also in the stratum radiatum (SR) and stratum lucidum (SL) of CA3 (Figure 3c,d). Finally, the association of Kv_1_ with the myelin proteolipid protein (MYPR), a major determinant in the formation and maintenance of the multilamellar structure of myelin is also lost in *Lgi1^-/-^*. All these molecular signatures indicate a drastic change in Kv_1_ membrane and sub-membrane associated complexes and corroborate the significant downregulation of membranous Kv_1_.^[8]^

**Figure 3.**
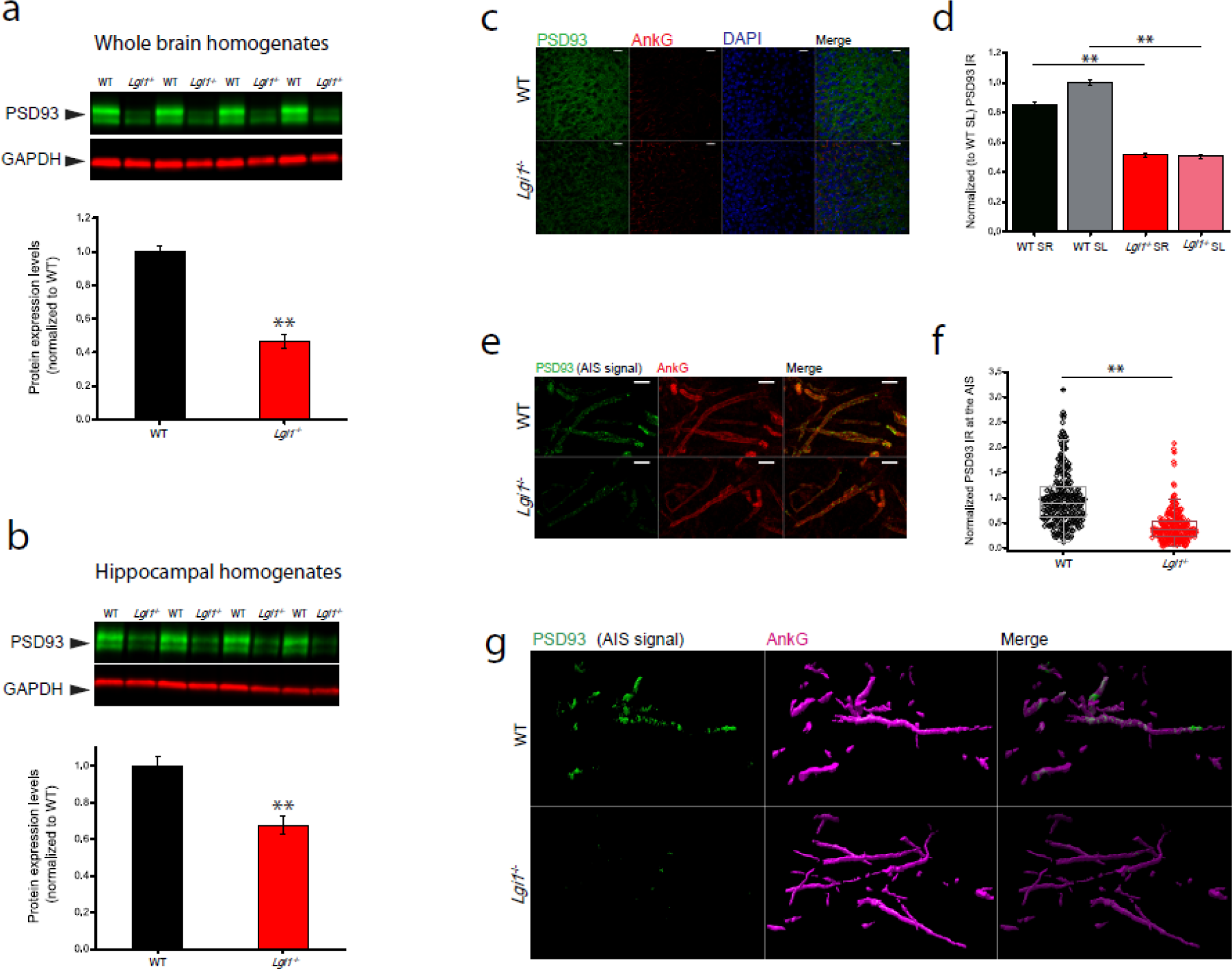
PSD93 expression is reduced in the *Lgi1*^-/-^ mice. a) Quantification by Western blot of PSD93 expression in brain homogenates of PSD93 in WT (1.00 ± 0.03) and *Lgi1*^-/-^ (0.46 ± 0.04); p = 1.40 x 10^-11^; n = 20; Two-sample t-test). b) Quantification by Western blot of PSD93 expression in hippocampal homogenates in WT (1.00 ± 0.05) and *Lgi1*^-/-^ (0.68 ± 0.05); p = 6.10 x 10^-5^; n = 20; Two-sample t-test). In a and b, signals were normalized to GAPDH expression levels. c) 25x representative images of Hippocampal CA3 region showing a decrease in PSD93 expression. PSD93 is in green, AnkyrinG is in red, and DAPI is in blue. Scale bar = 25µm. d) Quantification of PSD93 expression decrease across different CA3 regions in WT (black tones) and *Lgi1*^-/-^ (red tones) normalized to WT-SL (CA3-SR WT=0.85 ± 0.02 vs *Lgi1*^-/-^ = 0.51 ± 0.01; p = 9.65 x 10^-34^; CA3-SR WT = 1.00 ± 0.02 vs *Lgi1*^-/-^ = 0.50 ± 0.01; p = 1.31 x 10^-63^; ANOVA followed by Bonferroni’s test for means comparison). e) 63x representative images showing a reduction in PSD93 expression at the AIS level of CA3 Pyramidal neurons in *Lgi1*^-/-^; PSD93 is in green and AnkyrinG is in red; Scale bar=5 µm. f) Quantification of the decreased PSD93 expression at the AIS level in *Lgi1*^-/-^ hippocampal CA3 pyramidal neurons (WT = WT 1.00 ± 0.03 vs Kv1.1 *Lgi1*^-/-^ 0.42 ± 0.02, p = 0; Mann-Whitney U test). g) Surface representation of PSD93 labelling at the AIS of CA3 pyramidal neurons in WT (top panels) and *Lgi1^-/-^* (bottom panels) hippocampi. Magenta: AnkG, Green: PSD93.

#### 2.3.2. New Kv_1_ partners in Lgi1^-/-^ samples

*In parallel to* the loss of all the above mentioned membranous, sub-membranous and cytoskeletal links of Kv_1_, we found that, in *Lgi1^-/-^*(Table S3), Kv_1_ engages novel molecular interactions and shows association with the microtubule associated protein doublecortin (DCX) and with the GTPases Septin 6 and 7 that are involved in actin and microtubule cytoskeleton organization, membrane trafficking, vesicle transport and exocytosis as well as the assembly of scaffolding platforms.^[49 50]^ Interestingly, Septin 6 has been shown to be localized at the neck of dendrites ^[51]^, suggesting a potential reorganisation of Kv_1_ localization. In addition, we found that in *Lgi1^-/-^* samples, Kv_1_ is associated with the calcium sensor Copine 6 that regulates structural plasticity of spines. Despite the loss of interaction with actin, Kv_1_ in *Lgi1^-/-^* acquires association with the filamentous protein and actin-crosslinking MARCS (Myristoylated alanine-rich C-kinase substrate).

The acquirement of Kv1 association with VAMP2, syntaxin1A, synaptogyrin 3, Copine1, synapsin as well as dynamin 3 (DYN3) indicate the presence of Kv_1_ in vesicular compartments and suggest an increase in channel endocytosis. This is corroborated by the fact that neuronal activity blockade was shown to induce synaptic clustering of post-synaptic DYN3 ^[52 53 54]^ and that TTX-induced neuronal activity blockade induces a reduction of the Kv1-mediated DTX-I sensitive current.^55^ Also, it has been shown that activity-dependent Kv_1.2_ endocytosis from distal apical dendrites is mediated by dynamin3 in CA3 pyramidal.^[56]^ Therefore, the acquirement of DYN3 association with Kv_1_ may be the read out of this mechanism. Since Kv1 levels are reduced in *Lgi1*^-/-^, the association of Kv_1_ with DYN3 might reflect a mechanism of Kv_1_ retrieval at the dendritic level. This hypothesis is corroborated by the acquired Kv_1_ association in *Lgi1*^-/-^ with annexin A2 (ANXA2) which is a membrane-associated pleiotropic calcium-dependent phospholipid binding protein involved in several aspects of cell biology related to calcium signalling and endocytosis. ANXA2 is enriched in lipid rafts ^[57]^ and its secretion is activity-dependent.^[58]^ It has been shown to be primarily present in GABAergic interneurons ^[58]^ and is overexpressed in pathological neuronal and glial reactions.^[59]^ Therefore, its presence in the Kv_1_ interactome in *Lgi1^-/-^* background may be linked to the observed astrogliosis in these animals.

In addition to the perturbation of Kv_1_ cytoskeletal links, Kv_1_ transit from ER/Golgi to plasma membranes seems also to be affected since the association of Kv_1_ with the Golgi adaptors AP3S1 and AP1B1 ^[60]^ is disrupted. These adaptor proteins are known to play a crucial role in sorting the cargo destined to plasma membranes, neurites and nerve terminals.^[61 62]^

An ER population of Kv_1_ seems to be particularly prominent in *Lgi1^-/-^* samples. This conclusion is driven by the fact that in *Lgi1^-/-^*, Kv_1_ is associated with LRRC59 and RPN1. LRRC59 is an ER mRNA receptor known to participate in coupling translation and ER membrane insertion of transmembrane-domain containing proteins ^[63]^ and RPN1 (dolichyl-diphosphooligosaccharide-protein glycosyltransferase subunit 1 is involved in early glycosylation steps of nascent proteins. The association with the receptor of activated protein C kinase 1 (RACK1), PKCγ (KPCG), the cAMP-dependent protein kinase catalytic subunit beta (KAPCB) and the adenylate cyclase activator LANC2 ^[64]^ may reflect a change in phosphorylation-mediated dynamic balance and stability of Kv_1_ membrane expression. ^[65]^ Finally, the acquisition of association with L-lactate dehydrogenase B chain (LDHB) may indicate a functional modulation of channel activity since LDBH is part of the oxydoreductase complex and is involved in NAD metabolism which is important in Kv_1.2_ activity modulation.^[66]^ In parallel to this association, Kv_1_ coimmunoprecipitated with the voltage-dependent anion-selective channel protein 1 (VDAC1) that binds sphingolipids, cholesterol and phosphatidylcholine and is involved in apoptosis and energy metabolism.

#### 2.3.3. Significant changes in the association of some Kv_1_ partners in Lgi1^-/-^ samples

Among Kv_1_ partners that were immunoprecipitated from both wild type and *Lgi1^-/-^* extracts, those that varied less than ± 20 % were considered stable. ADAM22 (Disintegrin and metalloproteinase domain-containing protein 22) association was by far the most affected partner with over 60% decrease followed by DLG4 (PSD95) > DLG1 (SAP97) > DLG3 (SAP102) with >50 %, > 44% and 40% decrease respectively. Although both ADAM22 and ADAM23 co-immunoprecipitated Kv_1.2_ from pre-enriched LGI1-associated complexes ^[67]^, ADAM23 was totally absent from our immunoprecipitations from both WT and *Lgi1^-/-^* samples. Previous immunoprecipitations using anti-C-terminal Kv_1.2_ antibodies also did not show association with ADAM23.^[14]^ Whether the absence of ADAM23 is the result of epitope accessibility in Kv_1_ / ADAM23 complexes or reflects a real molecular organisation is still to be addressed.

Beside these partners that show decreased partnership with Kv_1_, an important increase in channel association with several key proteins was observed. The biggest increase in interaction (> 127%) is observed for 14-3-3γ (Ywhag). This increase is accompanied by a > 55% augmentation of interaction with 14-3-3ε. The second important increase (> 77%) concerned the vesicle-associated membrane protein-associated protein A (VAPA) but not VAPB that remains rather stable. VAPA/B are tether proteins of membrane contact sites between the endoplasmic reticulum (ER) and the plasma membrane (PM) ^[68]^ rich in AKAPs, kinases, Rabs and non-vesicular lipid transfer proteins (i.e ASTRA, ASTRB ^[69]^). VAPA/B possess a large number of common interactors, however, a specific interactome of each VAP exists yet without a clear functional discrepancy significance.^[68]^ The reason for the selective change in VAPA remains obscure, however, it may be related to the observed increase in the ER population of Kv_1_ and suggests that LGI1 is involved in the presence of Kv_1_ channels at ER/PM contact sites. Whether the increase in VAPA levels is linked to the increase in Kv_1_ association with ER proteins needs further investigations. In parallel to these differences, there was a significant increase of association with the cytosol aminopeptidase AMPL and the leucine-rich repeat and transmembrane domain-containing protein 2 (LRTM2), which are involved in axon guidance. It is interesting to note that while the interaction of Kv_1_ with PP2BB is lost, the one with PP2BA increases by >34%. In parallel to the interaction with Septin 6 and 7 mentioned earlier, an increase of interaction with Septin 11 was observed. Septin 11 (SEP11), is enriched post-synaptically at the neck of dendritic spines at GABAergic synapses. ^[70]^ This observation suggests a reorganisation in the dendritic localisation of Kv_1_. Among the partners with increased association with Kv_1_, we found WD repeat-containing protein 26 (WDR26) which is a G-beta-like negative regulator of MAPK pathway, which haploinsufficiency induces among other symptoms intellectual disability and seizures.^[71]^ Interestingly, the LGI1 EPTP domain shows sequence homology with WD40 sequences ^[25]^ and may reflect a particular propensity of Kv1 subunits to interact with some WD40 motif-containing proteins. This finding supports the above mentioned suggestion for the existence of changes in the phosphorylation-mediated dynamic balance and stability of Kv_1_ membrane expression ^[65]^ in *Lgi1^-/-^*. In parallel to the acquired association with endocytotic markers mentioned earlier, the association of Kv_1_ with syntaxin1B was also increased.

### 2.4. Kv_2_ expression is decreased in *Lgi1^-/-^*

Homo-tetrameric Kv_2_ delayed-rectifier potassium channels are widely expressed though hetero-oligomers can occur in some cells. ^[72]^ VAPA/B interact with Kv_2.1_ and Kv_2.2_ channel subunits ^[73]^ and induce the formation of channel clusters. ^[72]^ At PM/ER contact sites, Kv_2_ clusters constitute trafficking hubs where membrane protein insertion and retrieval take place. ^[74]^ As we observed an increase of VAPA association with Kv_1_ in *Lgi1^-/-^* and a loss of Kv_1_ association with the VAPA interaction partner ASTRA, we undertook to investigate whether Kv_2_ expression in *Lgi1^-/-^*was modified. As shown in Figure 4a and b, Kv_2.1_ expression was specifically and drastically reduced in hippocampus (Kv_2.1_ WT 1.00 ± 0.05 vs Kv_2.1_ *Lgi1*^-/-^ 0.45 ± 0.04, p = 1.00 x 10^-9^) compared to total brain homogenates (Kv_2.1_ WT 1.00 ± 0.09 vs Kv_2.1_ *Lgi1*^-/-^ 1.01 ± 0.12, p = 0.97) where no difference was observed. Kv_2_ channels provide a major component of somatodendritic potassium currents and occur at the neuronal surface either in conducting freely-diffusing forms or non-conducting clusters.^[75]^ These channels are involved in the dynamic modulation of excitability during periods of high neuronal activity.^[75 76]^ In addition, a decrease in Kv_2.1_ expression leads to an increase in the propensity of neurons to exhibit complex spike bursting. ^[29]^ In order to get more insight into potential modification in the expression of Kv_2.1_ in the *Lgi1^-/-^* hippocampal formation, we performed immunohistochemical staining of Kv_2.1_ in hippocampal slices. As described earlier ^[77]^, Kv_2.1_ was mostly enriched in the CA1 compared to the CA3 (Figure 4c). We found that in *Lgi1^-/-^,* the expression level of Kv_2.1_ in the CA3 was severely decreased (Figure 4c,e). A closer look at the distribution of this channel in the CA3 shows also considerable decrease of expression in the stratum oriens (SO) and stratum pyramidale (SP) (Figure 4d,f) and that AIS labelling was severely diminished (Figure 4f-h). In order to get more insight into the nature of the decrease in Kv_2.1_ staining, we analysed the following four distinct staining parameters, either in global staining or at the AIS: i) cluster density ii) normalized immunoreactivity of the clusters iii) cluster surface and iv) cluster volume. As shown in Figure 4e and Figure 4h, cluster density was the only staining parameter that showed significant decrease in the global analysis or at the AIS. This fully converges with the previously reported dispersion of Kv_2.1_ clusters in response to increased neuronal activity.^[44 78]^

**Figure 4.**
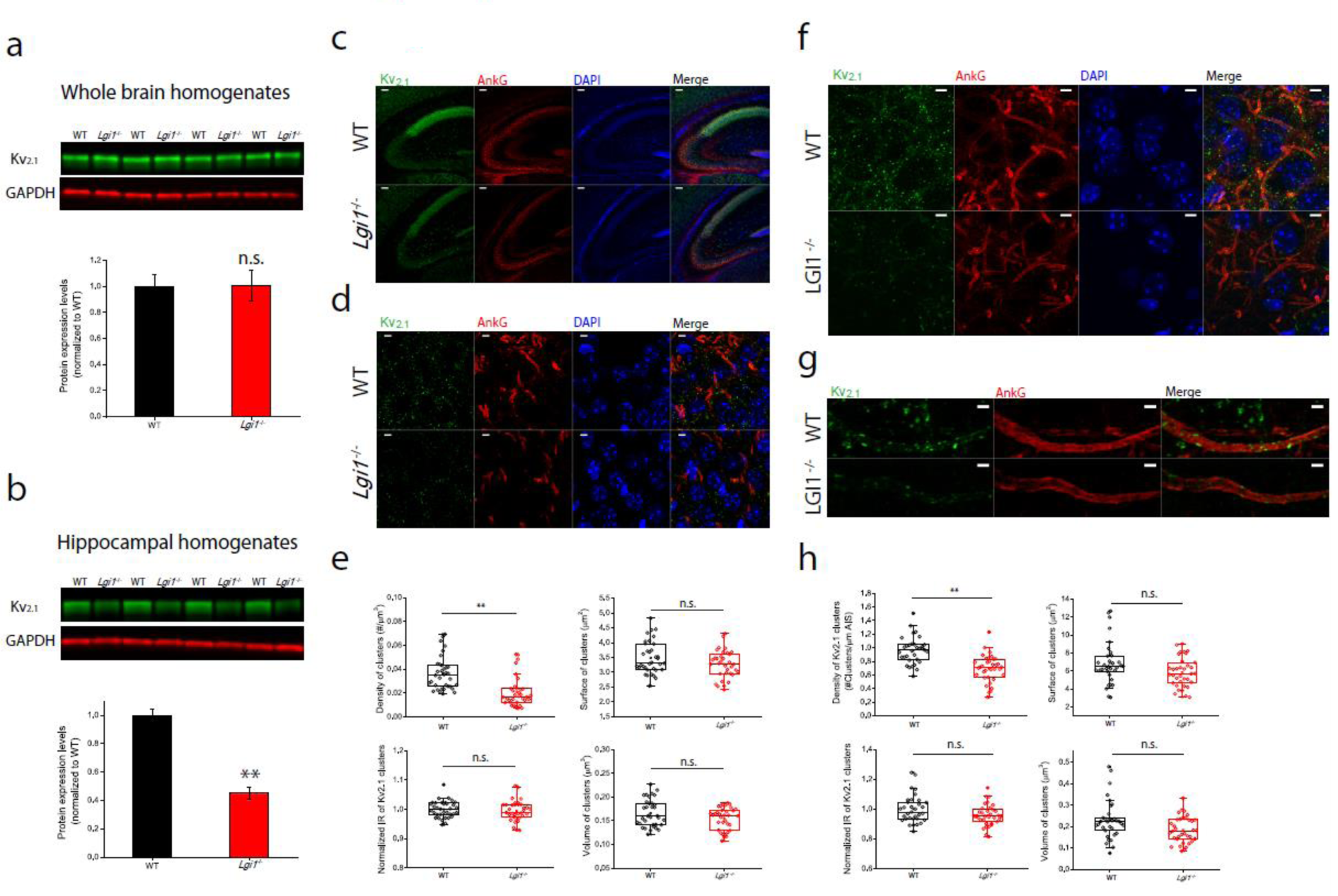
Kv_2.1_ is selectively reduced in hippocampus of *Lgi1*^-/-^ mice. a) Quantification by Western blot of Kv_2.1_ expression in brain homogenates of WT (WT 1.00 ± 0.09) and *Lgi1*^-/-^ (1.01 ± 0.11); p = 0.97; n = 15; Two-sample t-test). b) Quantification by Western blot of Kv_2.1_ expression in WT (1.00 ± 0.04) and *Lgi1*^-/-^ (0.45 ± 0.04) hippocampal extracts; p = 1.00 x 10^-9^; n = 16; Two-sample t-test). c) 10x representative images showing a reduced expression of Kv_2.1_ across the hippocampus of *Lgi1*^-/-^ mouse. Scale bar = 100 µm. d) Representative 63x images showing a reduction of Kv_2.1_ in CA3 Str. Pyramidale of *Lgi1*^-/-^ hippocampus. Scale bar = 5 µm. e) Analysis of Kv_2.1_ puncta in CA3 in WT (black) and *Lgi1*^-/-^ (red) hippocampi showing a reduction of Kv_2.1_ clusters’ density in *Lgi1*^-/-^. (Density of Clusters (#/µm^3^) WT = 0.036 ± 0.002 *Lgi1*^-/-^ = 0.020 ± 0.002; p = 1.53×10^-6^; Mann-Whitney U test; Surface of clusters (µm^2^) WT = 3.48 ± 0.09 *Lgi1*^-/-^ = 3.27 ± 0.08; p = 0.08; Two sample t-test; Normalized (to WT) immunoreactivity of clusters WT = 1.00 ± 0.01 *Lgi1*^-/-^ = 0.99 ± 0.01; p = 0.53; Two sample t-test; Volume of clusters (µm^3^) WT = 0.16 ± 0.01 *Lgi1*^-/-^ = 0.15 ± 0.01; p = 0.07; Two sample t-test). f) Representative 63x images showing a reduction of Kv_2.1_ at the level of CA3 Pyr. neurons AIS (CA3 Str. Oriens) of *Lgi1*^-/-^ hippocampus. Scale bar = 5 µm. g) 63 Detail of a CA3 Pyr. neuron AIS in WT (top) and *Lgi1^-/-^* (bottom) showing a reduced density of Kv_2.1_ clusters at the AIS of *Lgi1*^-/-^ neurons. Scale bar = 2 µm. h) Analysis of Kv_2.1_ puncta in the AIS of CA3 pyramidal neurons in WT (black) and *Lgi1*^-/-^ (red) showing a reduction of Kv_2.1_ clusters’ density and volume at the AIS of CA3 pyramidal neurons in *Lgi1*^-/-^ mouse. (Density of Clusters (#/µm AIS) WT = 0.96 ± 0.03 *Lgi1*^-/-^ = 0.69 ± 0.03; p = 4.58 x 10^-7^; Two sample t-test; Surface of clusters (µm^2^) WT = 6.87 ± 0.41 *Lgi1*^-/-^ = 5.85 ± 0.27; p = 0.07; Mann-Whitney U test; Normalized (to WT) immunoreactivity of clusters WT= 1.00 ± 0.02 *Lgi1*^-/-^ = 0.96 ± 0.01; p = 0.09; Mann-Whitney U test; Volume of clusters (µm^3^) WT = 0.23 ± 0.01 *Lgi1*^-/-^ = 0.18 ± 0.01; p = 0.046; Mann-Whitney U test).

### 2.5. Functional correlate of changes in Kv_1_ & Kv_2_ expression levels and interactome changes

We and others have previously shown that LGI1 is particularly enriched in the CA3 hippocampal subregions ^[79 80 23]^ and that it expression is crucial in controlling neuronal excitability in CA3 pyramidal neurons.^[8 81 82]^ Moreover, LGI1-dependent autoimmune limbic encephalitis is coincident with focal CA3 hippocampal atrophy in human patients ^[83]^. It has also been pointed out that a decrease in somatodendritic D-type potassium currents ^[84 85 86 87 88]^ and a weak expression of Kv_2_ channels ^[29]^ could promote the appearance of bursting patterns in hippocampal neurons. Although we have previously characterized the intrinsic excitability in CA3 pyramidal neurons in cultured hippocampal slices from *Lgi1^-/-^*mutant mice ^[8]^, it has been shown that cultured hippocampal slices develop, with time, spontaneous epileptiform-like activity.^[89]^ In order to examine LGI1 deficiency-mediated modification in excitability independently from potential artefacts induced by slice culture conditions, we examined intrinsic excitability of CA3 pyramidal neurons from acute slices. To examine changes in excitability at the CA3 hippocampal level, we performed current clamp recordings from CA3 pyramidal neurons of WT and their *Lgi1^-/-^* littermates (Figure 5a,b). Similar to organotypic slices, we observed a marked increase of the excitability in CA3 pyramidal neurons from *Lgi1^-/-^* defined by a pronounced decrease in the rheobase (WT = 119 ± 20 pA vs *Lgi1^-/-^* = 45 ± 4 pA; p = 5×10^-6^, Mann-Whitney U test) and a significant reduction of the latency to the first spike at the rheobase current (WT = 482 ± 40 ms vs. *Lgi1^-/-^* = 341 ± 30 ms; p = 0.007, Mann-Whitney U test) (Figure 5c, d,e). Neither the AP half-width (WT = 1.22 ± 0.02 ms vs. *Lgi1^-/-^* = 1.20 ± 0.02 ms; p = 0.55, Two samples t-test), nor amplitude (WT = 90.10 ± 0.93 mV vs. *Lgi1^-/-^* = 92.61 ± 1.00 mV; p = 0.07, Two samples t-test) were modified (Figure 5f-h). This is consistent with previous observations showing that the broadening of AP due to the loss or pharmacological inhibition of Kv_1.1_ is only noticeable in presynaptic but not in somatic APs.^[90]^ Interestingly, we observed a proportion of CA3 pyramidal neurons that exhibit spike bursts at the rheobase (Figure 5i,j). Although this kind of firing pattern has been previously described ^[91 29]^ in WT cells, the proportion of neurons exhibiting bursts was substantially larger in *Lgi1^-/-^*mice compared to their WT littermates (WT_bursting cells_ = 6.67%, [2/30 cells] vs. *Lgi1^-/-^* _bursting cells_ = 48.57%, [17/35 cells]; p = 0.0003 Fisher’s exact test) (Figure 5k,l). In order to address bursting propensity and based on a previous work that focused on this type of firing pattern in CA3 pyramidal neurons ^[29]^, we performed a different set of experiments in which neurons were challenged with either 300 pA or 600 pA current injection pulses (0.5ms). The proportion of cells bursting at 300pA was significantly larger in *Lgi1^-/-^* mice (WT = 12.5% [1/8 cells] vs. *Lgi1^-/-^* 87.5% [7/8 cells]; p = 0.01 Fisher’s exact test) and virtually all the *Lgi1^-/-^* cells challenged with 600 pA current injections exhibited a bursting pattern (WT = 12.5% [1/8 cells] vs. *Lgi1^-/-^* 100% [8/8 cells]; p = 0.001 Fisher’s exact test) (Figure 5m-o).

**Figure 5.**
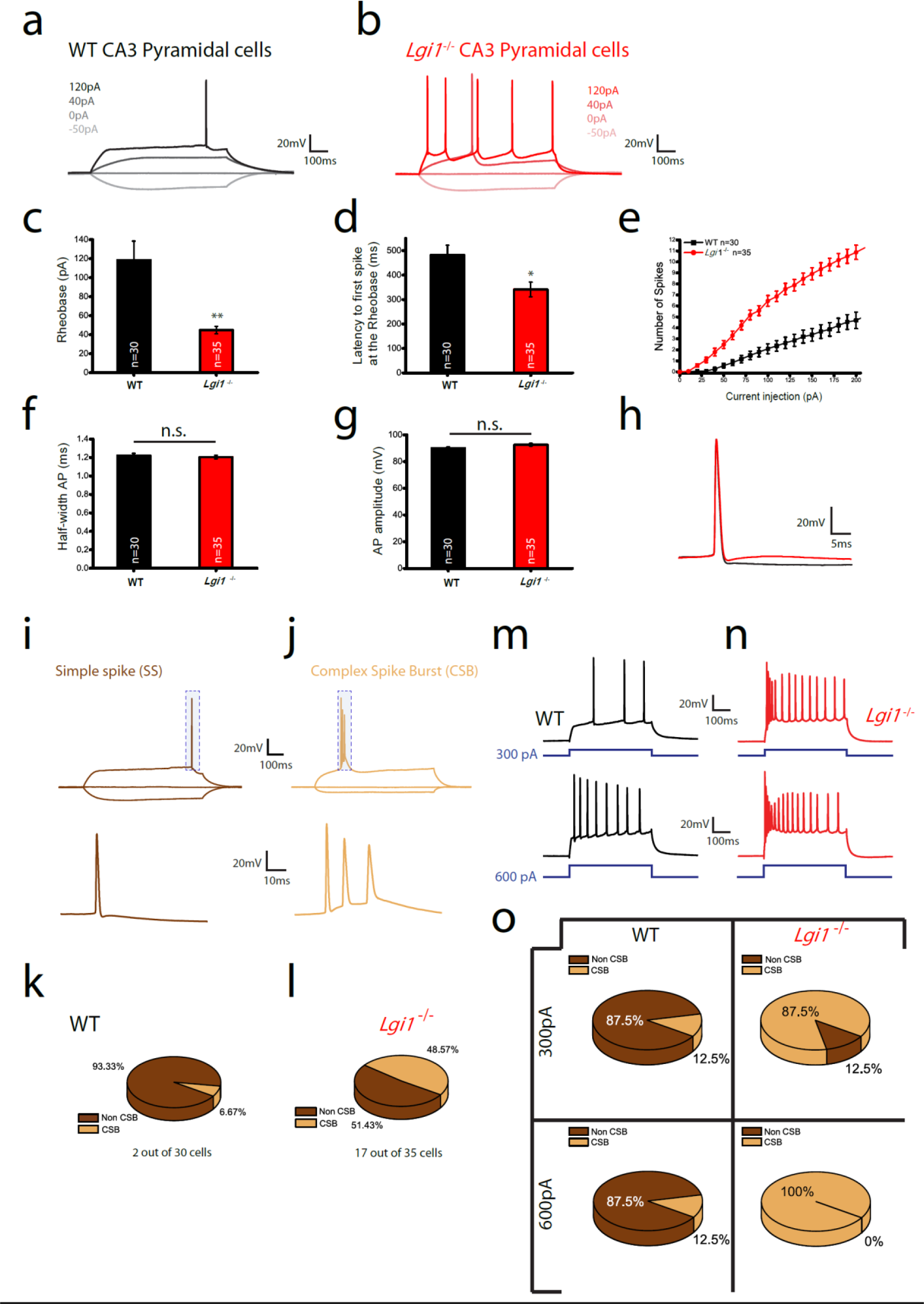
Intrinsic excitability and bursting propensity are increased in *Lgi1*^-/-^ CA3 Pyramidal neurons. a) representative traces for WT CA3 pyramidal neurons in response to the indicated current injections. b) representative traces for *Lgi1*^-/-^ CA3 pyramidal neurons in response to the indicated current injections. c) Average rheobase values for WT (black) and *Lgi1*^-/-^ (red) CA3 pyramidal neurons (WT = 116.00 ± 19.35 pA *Lgi1*^-/-^ = 44.86 ± 4.00 pA; p = 0.001; Two sample t-test). d) Average latency values to the first spike at the Rheobase in WT (black) and *Lgi1*^-/-^ (red) CA3 pyramidal neurons (WT = 475.60 ± 39.33 ms *Lgi1*^-/-^ = 341.40 ± 29.83 ms; p = 0.007; Two sample t-test). e) Average input-output curves of WT (black) and *Lgi1*^-/-^ (red) CA3 pyramidal neurons. f) Half-width of action potentials in WT (black) and *Lgi1*^-/-^ (red) CA3 pyramidal neurons (WT = 1.22 ± 0.02 ms *Lgi1*^-/-^ = 1.20 ± 0.02 ms; p = 0.55; Two sample t-test). g) Amplitude of action potentials in WT (black) and *Lgi1*^-/-^ (red) CA3 pyramidal neurons (WT = 90.09 ± 0.93 mV *Lgi1*^-/-^ = 92.61 ± 1.00 mV; p = 0.07; Two sample t-test). h) representative traces in current-clamp recordings showing the shape of an AP in WT (black) and *Lgi1*^-/-^ (red) CA3 Pyramidal neurons. i) Non-complex spike bursts (single AP, dark brown) and j) Complex-spike burst (light brown) recorded in current clamp in CA3 pyramidal neurons. k) proportion of CSB and Non-CSB neurons at the rheobase in WT CA3 pyramidal neurons. l) Proportion of CSB and Non-CSB neurons at the rheobase in *Lgi1*^-/-^ CA3 pyramidal neurons. m) Representative traces of bursting propensity in WT CA3 pyramidal neurons studied by suprathreshold current injections at 300 pA (top) and 600 pA (bottom) respectively. n) Representative traces of bursting propensity in *Lgi1*^-/-^ CA3 pyramidal neurons studied by suprathreshold current injections at 300 pA (top) and 600 pA (bottom) respectively. o) The proportion of CA3 pyramidal cells exhibiting a bursting phenotype is increased in the *Lgi1*^-/-^ mouse (at 300 pA CSB WT = 12.5% [1/8 cells] vs. CSB *Lgi1^-/-^* 87.5% [7/8 cells]; p = 0.01 Fisher’s exact test; at 600 pA CSB WT = 12.5% [1/8 cells] vs. CSB *Lgi1^-/-^* 100% [8/8 cells]; *p = 0.001* Fisher’s exact test)

## 3. Discussion

In this study we combined immunofluorescence and mass spectrometry experiments to address modifications in Kv_1_ distribution and interactome in *Lgi1^-/-^*. It was previously reported that Kv_1_ channels undergo a severe downregulation in mice lacking LGI1 ^[8]^ as well as upon *in vivo* hippocampal infusion of LGI1 antibodies.^[21]^ In this study, we performed a detailed immunohistochemical analysis of Kv_1.1_ and Kv_1.2_ expression at the *Lgi1*^-/-^ hippocampus. The use of antigen retrieval techniques, a necessary condition to reveal Kv_1_ immunoreactivity at the AIS of hippocampal neurons ^[92]^, allowed an accurate and reliable analysis of modifications of the Kv_1_ expression pattern at the cellular and subcellular levels. To our knowledge, this is the first time that Kv_1_ expression at the AIS is evaluated in fixed brains of *Lgi1*^-/-^ mice. In addition to previous observations on Kv_1_ downregulation at the AIS, we uncovered an important overall decrease in all neuronal compartments. This finding reflects more adequately the massive decrease in Kv_1_ expression measured in Western blots since the reduction of Kv_1_ population in the AIS cannot account for the > 50 % overall decrease. The stable expression of Kv_1.1_ and Kv_1.2_ in PV+ interneurons is consistent with the finding that the electrophysiological properties of PV interneurons were unchanged in *Lgi1*^-/- [19]^, and that deleting LGI1 in PV+ interneurons did not influence seizure susceptibility.^[93]^

Mass spectrometry analysis of Kv_1_-associated complexes in *Lgi1*^-/-^ revealed intricate molecular changes in the Kv_1_ interactome. Besides the major impact on the Kv1 association with the cytoskeletal network, the significant modification in Kv_1_ association with 14-3-3 proteins may be an important readout of the perturbations associated with the absence of LGI1. 14-3-3 are homo /hetero multimeric cytosolic phospho-proteins that are highly expressed in brain and implicated in cell signalling pathways. 14-3-3γ mutations have been shown to be associated with several brain disorders such as epileptic encephalopathy ^[94 95]^ as well as febrile seizures.^[96]^ Also, 14-3-3γ plays a pivotal role in oligodendrocytic apoptosis and neuroinflammation.^[97]^ Recently, it was shown that ADAM22 phosphorylation (S_832_) stabilizes its interaction with 14-3-3 proteins (ε, ξ, θ, γ, β, and η) and protects it from endocytosis.^[98]^ Of note, the loss of interaction of 14-3-3 with non-phosphorylated ADAM22 does not influence ADAM22 association with LGI1, Kv_1.2_ or PSD95. It is therefore tempting to speculate that in *Lgi1^-/-^* mice, a homeostatic cell response aiming at stabilizing membrane Kv_1_ drives an increase in ADAM22 / 14-3-3 / Kv_1.2_ association. Further investigation of the Kv_1_ phospho-proteome is still necessary to ascertain these suggestions. An additional important perturbation was the sharp reduction in the expression level of PSD93 in whole-brain and hippocampal homogenates, with a prominent reduction at the AIS of CA3 pyramidal neurons. Although PSD93 is one of the major constituents of LGI1 ^[67 99]^ and ADAM22-associated complexes ^[20]^, it poorly associates with Kv_1.2_ in whole brain extracts.^[14]^ The observed specific loss of interaction with PSD93 is consistent with the previously published data showing that PSD93 but not PSD95 ^[28 14]^ nor ADAM22 ^[14]^ is important in Kv_1.1_ and Kv_1.2_ channel localisation and clustering at AIS.^[5]^ The selective decrease in PSD93 expression contrasts with the reported stability of PSD95 in the *Lgi1^-/-^* background ^[100]^ and shows that the association of Kv_1_ with PSD93 but not SAP97, SAP102 nor PSD95 is dependent on the presence of LGI1. Interestingly, RhoA activity has been shown to be significantly increased in *Lgi1^-/-^*.^[101]^ Also, RhoA activity mediates spine morphological and functional plasticity ^[102 103]^ and may therefore be linked to the observed actin cytoskeleton remodelling. Interestingly, PSD93 phosphorylation by RhoA was recently shown to decrease its binding to ADAM22 and to increase PSD93 heterodimerization with PSD95.^[99]^ The only partner of Kv_1_ that interacts with PSD95/PSD93 and that is specifically lost in our immunoprecipitations from *Lgi1^-/-^* tissue is the plasma membrane calcium-transporting ATPase 4 (AT2B4 / PMCA4) that was shown to be enriched in lipid-rafts.^[104]^ AT2B4 interaction with PSD95/93 has been shown to increase with neuronal excitability and glutamate release.^[105]^ This increase in excitability leads to the activation of NMDA receptors and triggers AT2B4 selective internalization.^[106]^ In addition to these observations, it is known that the activity of AT2B4 is inhibited by 14-3-3ε ^[107]^ which interacts with and modulate the expression of the LGI1-binding partner ADAM22. Despite that the protein expression level of AT2B4 did not show any significant difference between WT and *Lgi1^-/-^* samples (Figure S5 Supplementary Information) the loss of PSD93 co-immunoprecipitation with Kv_1_ in *Lgi1^-/-^* may partially reflect the loss of Kv_1_ association with AT2B4. Since LGI1 was shown to regulate PSD95-mediated AMPA and NMDA receptor levels in neurons ^[100]^, it may be suggested that, in the *Lgi1*^-/-^ epileptic model, a hyperexcitability-driven increase of interaction between PSD95 and AT2B4 could be blocked by the expected increase in RhoA-mediated PSD95/PSD93 interaction. Of note, it was recently reported that the truncation of the MAGUK binding domain of ADAM22 induces a massive reduction in hippocampal PSD95 expression levels.^[20]^ In line with these observations, the loss of PSD93 co-immunoprecipitation with Kv_1_ in *Lgi1^-/-^* may be the result of a RhoA-kinase phosphorylation of PSD93 and a concomitant delocalization of the PSD95/PSD93 complex. Further analysis is still necessary to understand the specificity of the seemingly key disruption of PSD93 association with Kv_1_ complexes in *Lgi1^-/-^*.

The most abundant partner of Kv_1_ that we find in *Lgi1^-/-^* is the glyceraldehyde-3-phosphate dehydrogenase (G3P) (Table S3, Figure S6) which is found with an emPAI (1.64 ± 0.34) close to the emPAI value of actin (2.12 ± 0.0) in wild type tissue (Table S2, Figure S6). G3P is a moonlighting protein involved in several regulatory pathways including synaptic transmission and intermembrane trafficking.^[108]^ The second most abundant acquired interaction partner is the fatty-acid binding protein FABP7 which is involved in the uptake of fatty acids and serves as a cellular chaperone of lipophilic molecules. This protein is expressed in astrocytes and oligodendrocyte progenitor cells (OPCs) ^[109 110]^ and has been shown to regulate dendritic morphology and excitatory synaptic function of cortical neurons.^[111]^ This observation is in line with the potential expression of Kv_1.2_ in astrocytes.^[112]^ Interestingly, among the newly acquired partners we found PCMD2 (protein-L-isoaspartate O-methyltransferase domain-containing protein 2) a member of the L-isoaspartyl O-methyltransferases (PIMT) protein family, known to repair natural protein damage.^[113 114 115 116]^ The presence of this protein in Kv_1_-associated complexes may suggest that at least some of its components may be under repair and imply a higher level of protein damage/recycling than in WT tissue. The presence of synapsin 2 (SYN2) and the voltage-dependent anion-selective channel protein 1 (VDAC1) in the Kv_1_ interactome of *Lgi1^-/-^* corroborates this hypothesis since these proteins have been identified as major endogenous substrates for PIMT.^[116]^ The association with the ATPase-dependent chaperone T-complex protein 1 subunit delta (TCPD) is in line with this interpretation and may highlight attempts for recovery from damage. This protein localises to dendrites and is involved in microtubule organisation in dendrites.^[117]^

Trafficking pathways associated with Kv_1_ also seem to have been affected in *Lgi1^-/-^* tissue, as the presence of AP1- and AP3-associated coat protein PACS1 decreased by >29% and the association with PSD1 (Cytohesin 1), a guanine nucleotide exchange factor for ARF6 involved in neuron projection development, protein trafficking and cytoskeletal rearrangements ^[118]^, decreased by > 38 %.

All these results underscore the complex nature of Kv_1_ interactions, where the absence of LGI1 leads to a complex rewiring of protein partnerships, resulting in both gains and losses that intricately modulate the network dynamics and underly a complex deleterious phenotype in the *Lgi1^-/-^* mouse.

In parallel to the direct observations driven by mass spectrometry analysis, we report for the first time a massive reduction in Kv_2_ in *Lgi1^-/-^.* In addition to the crucial role of Kv_1_ channels in tuning the excitability of CA3 pyramidal neurons ^[119]^, its decrease at the somatodendritic compartment may trigger an increased after-depolarization.^[84]^ Moreover, the reduction in Kv_2_ expression may cause an increase in dendritic calcium spikes. These two different mechanisms could act synergistically facilitating bursting.^[84] 29]^ Our results are also in agreement with other described forms of increased excitability in *Lgi1^-/-^* mice ^[120 19 121]^, and with the observed increase in excitability upon the treatment of slices with LGI1 antibodies.^[122 81 82]^ The importance of Kv_2_ downregulation in perturbing excitability in LGI1 deficiency is also highlighted by the fact that restoring Kv_1.1_ expression in secretion-deficient LGI1^W183R^ neurons does not fully rescue the mice life span.^[22]^ Further investigations are needed to address whether LGI1 is directly implicated in modulating Kv_2_ expression levels and whether Kv_1_ downregulation is parallel to or consequent to Kv_2_ downregulation.

In the light of the concomitant decrease in Kv_1_ and Kv_2_ expression levels, we addressed the intrinsic excitability properties of CA3 pyramidal neurons of *Lgi1^-/-^*. We and others have previously shown that LGI1 is particularly enriched in CA3 hippocampal subregion ^[79 80 23]^ and that LGI1 is crucial in controlling neuronal excitability in CA3 pyramidal neurons.^[8 81 123]^ Moreover, LGI1 antibody-mediated limbic encephalitis is coincident with focal CA3 hippocampal atrophy in human patients.^[83^ Although we have previously characterized the intrinsic excitability in pyramidal CA3 neurons of *Lgi1^-/-^* cultured hippocampal slices ^[8]^, it has been proposed that cultured hippocampal slices spontaneous develop epileptiform activity.^[89]^ To eliminate a potential bias introduced by the use of organotypic cultures, and to gain further understanding of the mechanisms underlying LGI1 deficiency-mediated epilepsy, we analysed intrinsic excitability of CA3 pyramidal neurons from acute slices.

We observed an increase in neuronal burst firing propensity and intrinsic excitability defined as a decrease in the rheobase and a shortened latency to the first spike. This gain in excitability converges with the decrease in Kv_1.1_ and Kv_1.2_ as well as Kv_2_ described throughout this study, and is in accordance with the crucial role of Kv_1_ channels in tuning CA3 pyramidal neurons excitability ^[119]^ and the implication of dendritic D-type potassium currents ^[84]^ and Kv_2_ channels ^[29]^ in the promotion of bursting patterns in hippocampal neurons. The presented results are also in agreement with other described forms of increased excitability in *Lgi1*^-/-^ mice ^[120 22 121]^, and with the observed increase in excitability upon the treatment of slices with LGI1 antibodies.^[122 81 123]^ Expectedly, there was no change in AP waveform since the broadening of the AP due to the loss or pharmacological inhibition of Kv_1.1_ is only noticeable in presynaptic, but not in somatic APs.^[90]^

Although these promising results deserve further investigation, it is widely accepted that synchronized bursting is the cellular substrate for interictal spikes, which, in turn, constitute a hallmark of epilepsy. ^[124 125]^ Given the high degree of recurrent connectivity in the CA3 hippocampal region ^[126 127]^, it is tempting to propose a cellular landscape in which the abnormally increased burst firing, in combination with a lower firing threshold in *Lgi1^-/-^* CA3 pyramidal neurons, triggers the epileptogenic phenotype observed in *Lgi1^-/-^* mice.

## 4. Experimental Section

### Antibodies and other reagents

Mouse (K20/78; RRID:AB_10672854 and K36/15; RRID:AB_10673166) anti-Kv_1.1_, anti-Kv_1.2_ (K14/16; RRID:AB_10674277) and anti-PSD93 (N18/28; 75-057, RRID: AB_2277296) were from Antibodies Incorporated and were used for Western blots and/or immunohistofluorescence experiments. Guinea pig (386005, RRID: AB_2737033) anti-ankyrinG (AnkG) was from Synaptic Systems and rabbit anti-Kv_2.1_ (APC-012) from Alomone labs. Monoclonal JA9 anti-PMCA4 was purchased from Thermo Fisher Scientific. Secondary Alexa coupled 488-goat anti-mouse, 488-anti-rabbit, 594-goat anti-guinea pig were from Jackson ImmunoResearch. A rabbit polyclonal anti C-terminal peptide (aa 454-468) of mouse Kv_1.2_ was produced and affinity purified by GeneCust and used for immunoprecipitation experiments. This antibody was used at 5 µg/ml for Western blot analysis and 5 µg / immunoprecipitated sample. HRP-coupled secondary antibodies were from Jackson ImmunoResearch. Chemicals were purchased form Euromedex and DAPI from Sigma-Aldrich.

### Clones

pcDNA3 plasmids containing human Kv_1.1_ or rat Kv_1.2_ were a generous gift from Jeffrey Martens (University of Michigan, USA). These sequences contain pHLuorin (Kv_1.1_) and EGFP (Kv_1.2_) coding sequences inserted in frame, in the extracellular loop linking transmembrane domains one and two. EGFP sequence in Kv_1.2_ was replaced by that encoding pHLuorin, using EcoR1 and Not1 restriction sites. The full coding sequences of pHLuorine-Kv_1.1_ and pHLuorine-Kv_1.2_ were transferred into pINDUCER11 (gift from Stephen Elledge, Harvard Medical School, Boston, USA) plasmid downstream of a doxycycline inducible promotor.

Kv_1.1_, Kv_1.2_, Kv_1.3_ and Kv_1.4_ C-terminal tail constructs were generated by standard polymerase chain reaction (PCR) using a commercial adult rat brain cDNA library (Origine). Sequences were cloned in pGEX-5X using EcoR I and Sal I restriction sites.

### Bacterial recombinant protein expression and purification

GST-Kv C-terminal tail fusion constructs were transfected into the BL21 bacterial strain and recombinant protein expression induced by 0.5 mM IPTG. Bacterial pellets were resuspended and solubilised in Tris 25 mM pH7.4, NaCl 150 mM supplemented with 1% Triton X-100 in the presence of protease inhibitors (Complete, Roche Diagnostics GmbH). Fusion proteins were purified on glutathione-Sepharose (Cytiva) using previously described procedures.^[128]^

### Western blots and ELISA

Western blots and ELISA experiments were performed using previously described procedures.^[129]^ For Western blots, 50 μg of proteins were loaded in each well. Signals were quantified using ImageJ and normalized to GAPDH intensities. For ELISA, 1 μg of purified GST-fusion proteins was immobilized in each well and proteins were detected using either anti-GST (1:2000) or anti-Kv_1.2_ polyclonal antibody. Anti Kv_1.2_ signals were normalized to anti-GST signals on parallel wells.

### Cell culture

HEK293T cells were cultured at 37°C in DMEM (GlutaMAX) supplemented with 10% foetal bovine serum, 1 % non-essential amino acids and 1% streptomycin/penicillin mix. Transfection with pINDUCER11-Kv_1.1_ or pINDUCER11-Kv_1.2_ expression plasmids was performed in optiMEM using Lipofectamine 2000. Kv expression was induced by doxycycline for 48h before cells harvesting. Cells were lysed in a Potter homogenizer using 20 mM HEPES pH7.4, NaCl 150 mM supplemented with protease inhibitors (Complete, Roche Diagnostics GmbH). Cell membranes were harvested by a 30 min centrifugation at 16000 x g and resuspended in lysis buffer for protein analysis.

### Biochemical sample preparations and immunoprecipitations

Wild type and *Lgi1^−/−^* ^[130]^ (the original *Lgi1^−/−^* mice strain was a generous gift from Stéphanie Baulac, ICM, Paris, France) mice total brains or hippocampi (15-16 days) were extracted, snap-frozen in liquid nitrogen and kept at -80°C until use. Homogenisation (1 ml / brain) was performed in HB buffer (20 mM Na phosphate pH 7.4, 30 mM NaCl containing phosphatase inhibitors (Pierce^TM^) and protease inhibitors (Complete, Roche Diagnostics GmbH) and nuclei were eliminated by 900 x g centrifugation. Supernatants were either directly processed for SDS-PAGE analysis and Western blot or solubilized for immunoprecipitation in HB supplemented with 1% CHAPS at 5 mg/ml and subjected to 100.000 x g ultracentrifugation. The solubilised material from both wildtype and *Lgi1*^-/-^ samples was precleared with rProtein A Sepharose Fast Flow (Cytiva) and immunoprecipitated in triplicates using the polyclonal anti-Kv_1.2_ antibody (3-5 µg / immunoprecipitated sample). Immunoprecipitated samples were washed three times with HB containing 0.5% CHAPS before denaturation. Samples were then analysed either by SDS-PAGE and Western blot or by mass spectrometry. C3HeB/FeJ Kcna2 mice brains (P10) (Kv ^-/-^) and their wild type littermates ^[56]^ were extracted and deep frozen in dry ice then transferred to -80°C and stored until use. Homogenisation and analysis by Western blot were performed as described for WT and *Lgi1^-/-^* samples.

### Mass spectrometry

Samples were treated and analyzed as in.^[23]^ Briefly, immunoprecipitated proteins were concentrated and analyzed by mass spectrometry using a hybrid Q-Orbitrap mass spectrometer (Q-Exactive, Thermo Fisher Scientific, United States) coupled to nanoliquid chromatography (LC) Dionex RSLC Ultimate 3000 system (Thermo Fisher Scientific, United States). All raw data files generated by MS were processed to generate mgf files and screened against the Swissprot database (Swissprot version june 2019, (560,118 sequence), with the taxonomy Mus musculus, using the MASCOT software (www.matrixscience.com, version 2.3). The database was searched in the decoy mode and emPAIs automatically calculated by the MASCOT algorithm. All samples were analysed in triplicates. For each antibody, background interactomes obtained with non-immune antibodies were subtracted using an Excel script. Keratins, ribosomal proteins, DNA binding proteins, mitochondrial proteins, transcriptional proteins and immunoglobulins were excluded from the list of candidates. Candidates with emPAI <0.1 were not interpreted. Functional properties of candidate proteins were classified using the pantherdb web site (www.pantherdb.org) (Supplementary sheet). The mass spectrometry proteomics data have been deposited to the ProteomeXchange Consortium via the PRIDE ^[131]^ partner repository with the dataset identifier PXD043649.

### Immunohistofluorescence staining of fixed brains

Fixed brains of P14-P16 C57BL/6 wild-type mice or *Lgi1*^-/-^ littermates of either sex were sliced and stained as described in.^[23]^ For Kv_1.1_ and Kv_1.2_ staining, an antigen retrieval process was applied as in.^[33 32]^

### Image analysis

image analysis was performed as described in.^[23]^ Briefly, AnkyrinG staining was used to measure the immunoreactivity of Kv_1_._1_, Kv_1_._2_, Kv_2.1_ and PSD93 ascribable to the AIS. For 3D quantification in confocal images, 3D objects counter ^[132]^ and 3D ROI manager ^[133]^ were integrated into custom-written FIJI scripts available upon reasonable request. In the sake of clarity in Figure 3e, the PSD93 signal corresponding to the AIS was extracted by using the “Clear Outside” method in ImageJ after using the thresholded AnkyrinG signal as a 3D selection.

### Acute Slice preparation

All experiments were performed in accordance with the European and institutional guidelines for the care and use of laboratory animals (Council Directive 86/609/EEC and French National Research Council) and approved by the local authority (Préfecture des Bouches-du-Rhône, Marseille). Mice of either sex were used in this study.

Postnatal (P14) WT or *Lgi1^-/-^* mice were deeply anesthetized in isoflurane (3% in oxygen, 2 L/min) and decapitated. After brain removal, the brains were immersed in oxygenated (95% O_2_, 5% CO_2_) ice-cold low-calcium artificial CSF (aCSF) containing the following (in mM): 125 NaCl; 2.5 KCl; 1.25 NaH_2_PO_4_; 0.5 CaCl_2_; 2 MgCl_2_; 26 NaHCO_3_; 10 D-Glucose; pH 7.4. Horizontal slices (350µm) were cut in a vibratome (Leica VT1200S) in oxygenated ice-cold low-calcium aCSF. Afterwards, slices were transferred and kept at least 1h at RT in oxygenated aCSF containing the following (in mM): 125 NaCl; 2.5 KCl; 0.8 NaH_2_PO_4_; 26 NaHCO_3_; 3 CaCl_2_; 2 MgCl_2_; 10 D-Glucose; pH 7.4.

### Electrophysiology and data analysis

Whole-cell recordings of visually identified CA3 pyramidal neurons were performed in a temperature controlled (30°C) chamber superfused with oxygenated aCSF. Borosilicate patch pipettes (3-5 MΩ) were pulled on a DMZ-Universal Puller (Zeitz Instruments) and filled with an internal solution containing (in mM): 120 K-gluconate; 20 KCl; 10 HEPES; 0.1 EGTA; MgCl_2_; 2 Na_2_ATP; 0.3 NaGTP; pH 7.4; 290-300 mOsm. Recordings were made in current clamp mode with a multiclamp 700B amplifier (Molecular Devices) and pClamp10 software. Recordings were acquired at 10 kHz and filtered at 3 kHz. Neurons with a resting membrane potential > -55mV were discarded from the final analysis. Intrinsic excitability was assessed by injecting a range of 0.8s current pulses (ranging from -50 to 250 pA;10 pA / step) and counting the number of spikes. Input–ouput curves were determined and rheobase was defined as the minimal current eliciting at least one action potential. To study bursting propensity, depolarizing current injections of 0.5s (300 or 600 pA) were applied every 2 s. Bursts were scored when ≥ 2 APs appeared in the same response showing progressively smaller amplitude and longer duration.^[29]^ To analyse the proportion of cells exhibiting complex bursts, only cells showing bursts at the rheobase current injection were considered. To study burst propensity, only cells exhibiting bursts when challenged with either 300 pA or 600 pA depolarizing pulses were defined as positive. AP parameters (AP half-width and AP amplitude) were extracted from the first spike in the rheobase. To prevent synaptic contamination 3mM kynurenate and 100µM picrotoxin (both from SIGMA) were systematically added to the aCSF. Cells were held at ∼ -70 mV during experiments and recordings were not corrected for liquid junction potential. All data were analyzed using ClampFit (Molecular Devices). Data are presented as mean ± SEM. Mann–Whitney U test, Two samples T-test, and Fisher’s exact test were used for comparisons. Normality tests were applied before the use of the Mann-Whitney U test or Two samples t-test.

## Acknowledgements

We thank Stephanie Baulac for sharing the *Lgi1^−/−^* mice strain ^[130]^ and Jeffrey Martens (University of Michigan, USA) for providing Kv1 clones ^[134]^.

## Author Contributions

O.E.F. conceived the study, supervised the entire project, the experimental design, data interpretation and manuscript preparation. O.E.F. and J.R-F. designed the study, analysed and interpreted the data. M.Se. and C.L. contributed to the design of the study. J.R-F. performed immunofluorescence experiments on slices, images treatments and analysis. J.R-F. performed electrophysiological recordings and Excel-based mass spectrometry analysis. K.D. performed Western blots, immunoprecipitations and participated in immunofluorescence experiments and data analysis. J.R-F. and K.D. took care of mice breeding and availability of *Lgi1^-/-^* animals. L.SH and Y.K. provided Kv_1.2_ brain tissues. M.B. performed mass spectrometry experiments. M.S. performed expression plasmids preparations and preliminary expression tests of recombinant constructs in heterologous systems. O.E-F. and J.R-F. wrote the original draft of the manuscript. O.E-F. and J.R-F. prepared the figures. All authors edited and reviewed the manuscript.

## Data Availability Statement

The data that support the findings of this study are available in the supplementary material of this article.

## Supporting information

**Figure S1.**
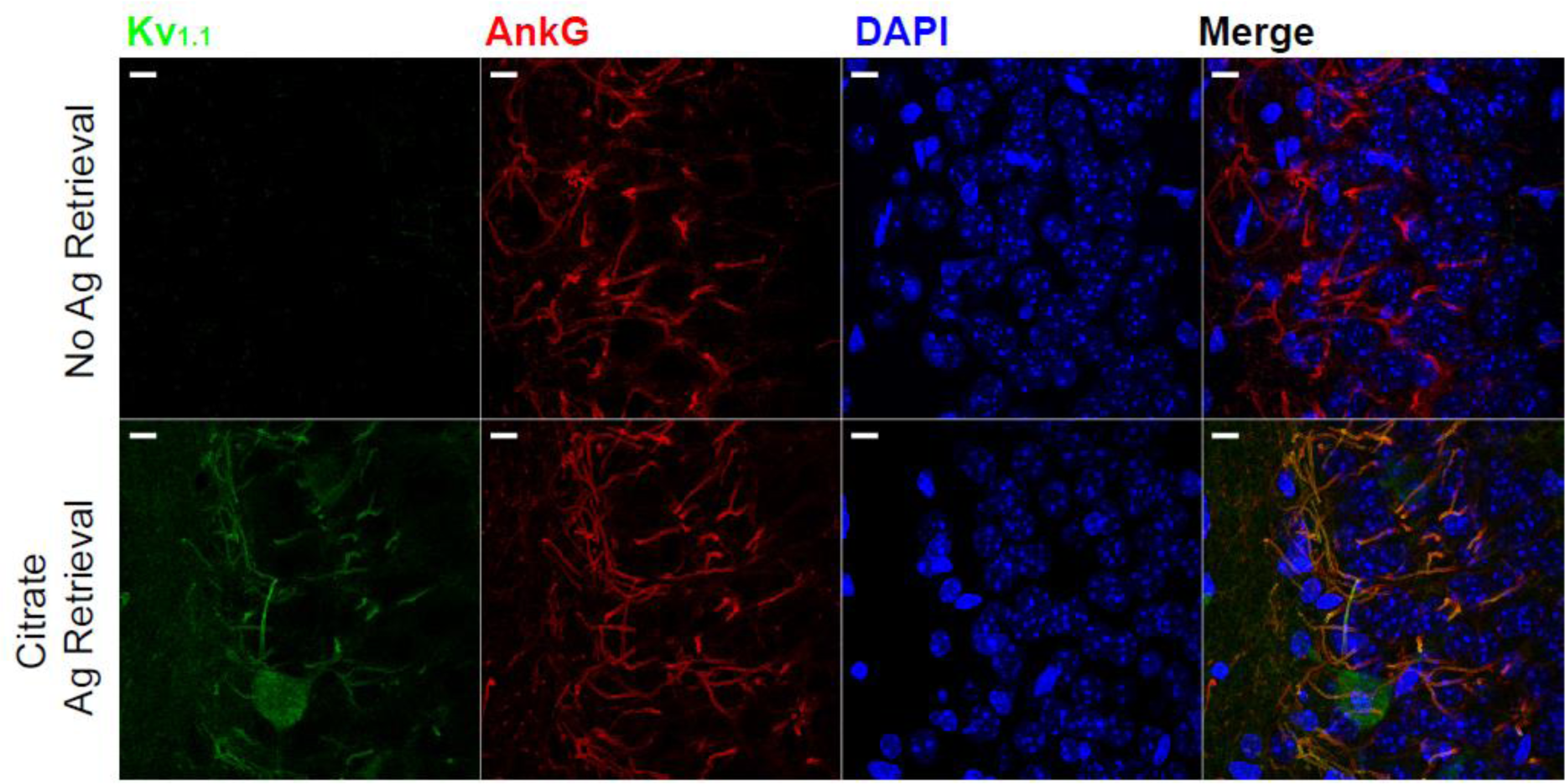
Antigen retrieval is necessary for specific Kv_1.2_ detection in brain slices. Fixed brains of P14-P16 C57BL/6 wild-type mice were sliced and probed with anti Kv_1.1_ and AnkG antibodies in the absence (upper panels) or presence (lower panels) of antigen retrieval. Note that AnKG staining was not different in both cases while Kv_1.1_ staining was only possible after antigen retrieval. Scale bar = 12.5 µm

**Figure S2.**
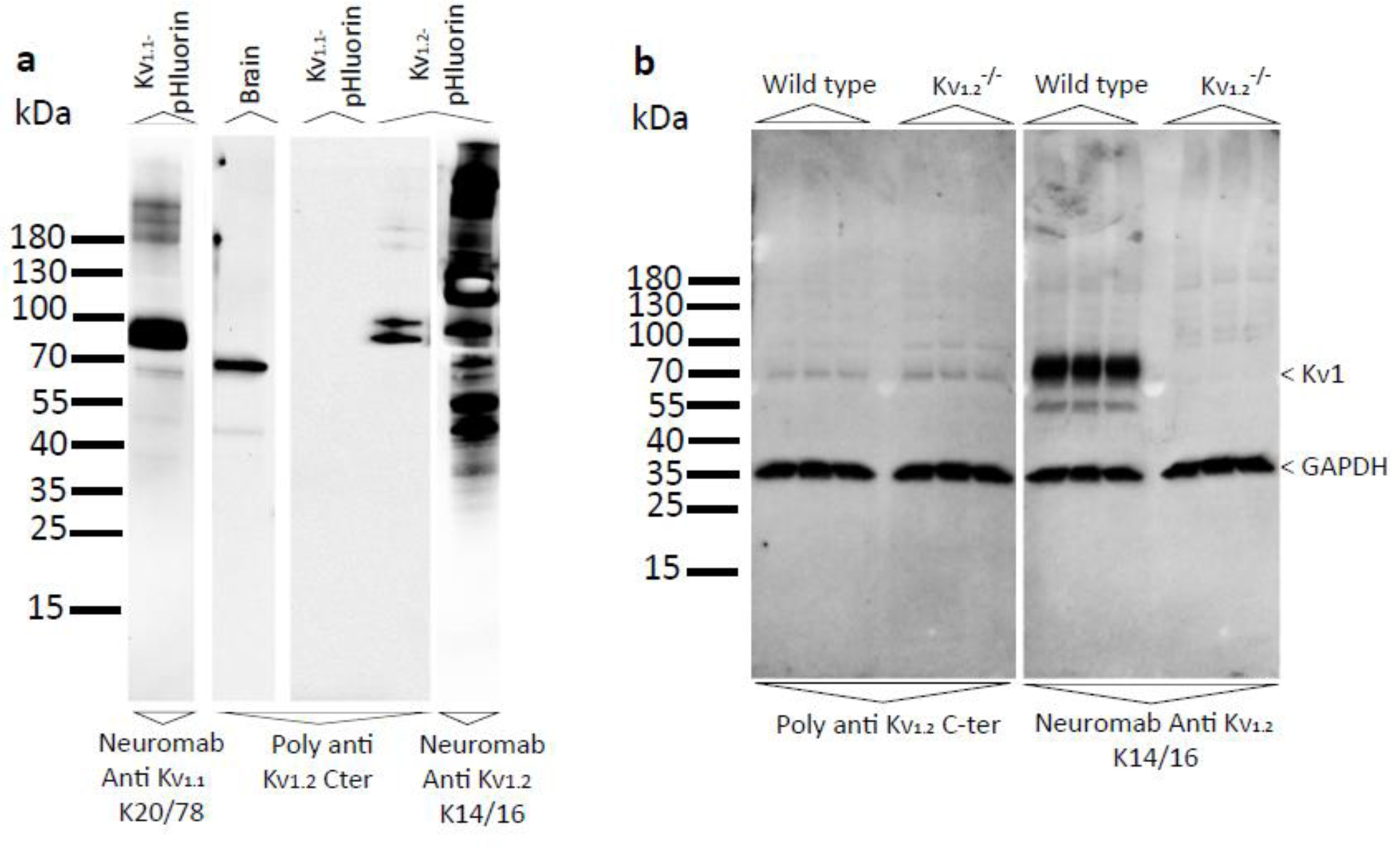
Characterisation of the rabbit polyclonal Kv_1.2_ antibody. HEK cell extracts transfected with recombinant Kv_1.1_ and Kv_1.2_ as well as brain homogenates were probed with the polyclonal Kv_1.2_ antibody. Control recombinant channels expression were performed using the Neuromab Kv_1.1_ and Kv_1.2_ antibodies. b) Western blots of wild type and Kv_1.2_^-/-^ brain homogenates with the rabbit polyclonal Kv_1.2_ antibody showing cross-reactivity with closely related native Kv1 channels.

**Figure S3.**
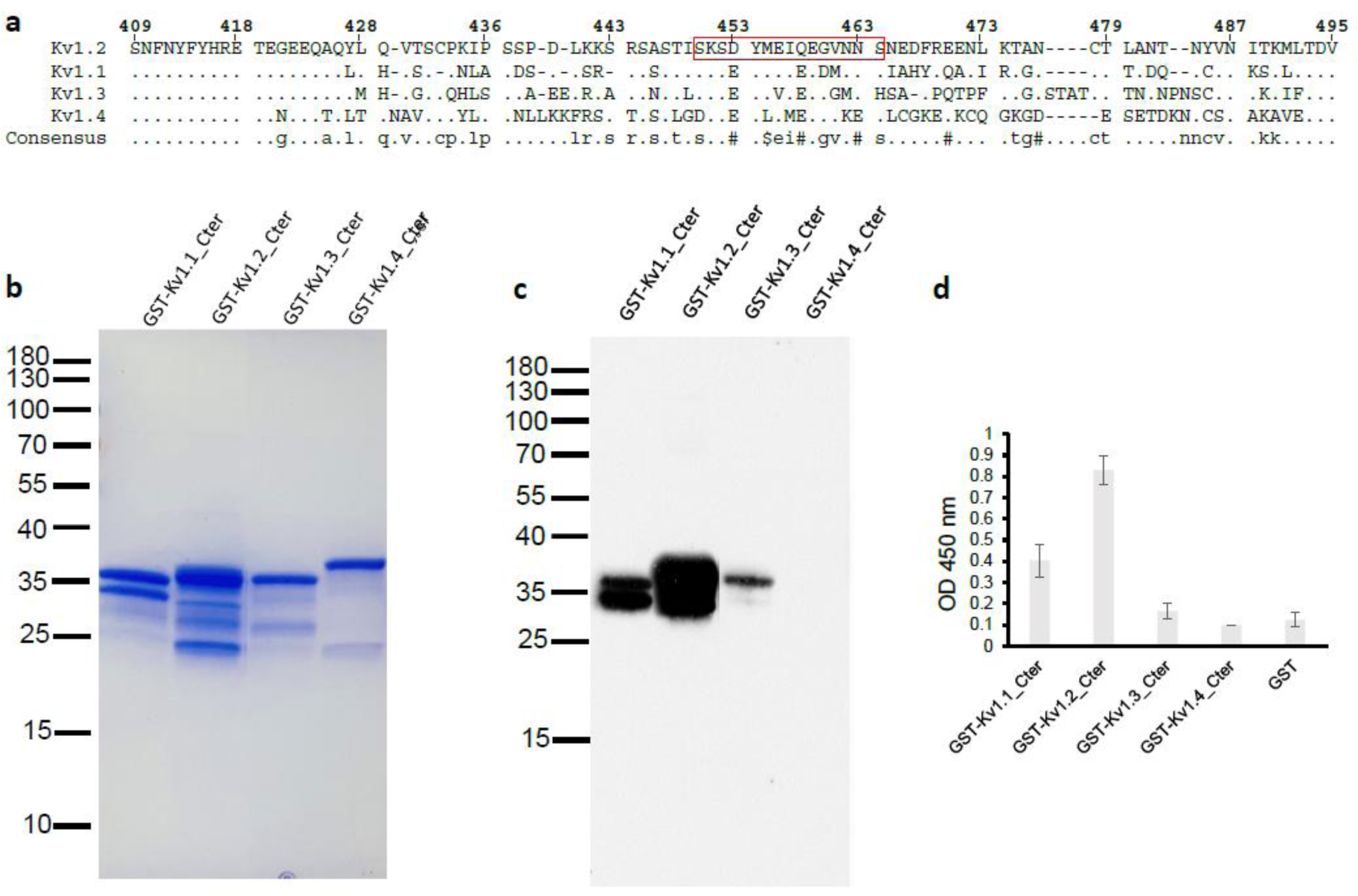
Isoform specificity of the rabbit polyclonal Kv_1.2_ antibody. a) Alignment of the amino acid sequences of C-terminal tails of mice Kv_1.1_, Kv_1.2_, Kv_1.3_, Kv_1.4_. The peptide sequence that served to generate the antibody is boxed. b) Coomassie staining of the different GST fusion protein constructs (3 μg) used for western blot. c) and d) Western blot (300 ng/well) and ELISA (0.5 μg/well) respectively of the constructs shown in b with the polyclonal Kv_1.2_ antibody showing a cross-reactivity of this antibody with Kv_1.1_ and to very slight extent with Kv_1.3_.

**Figure S4.**
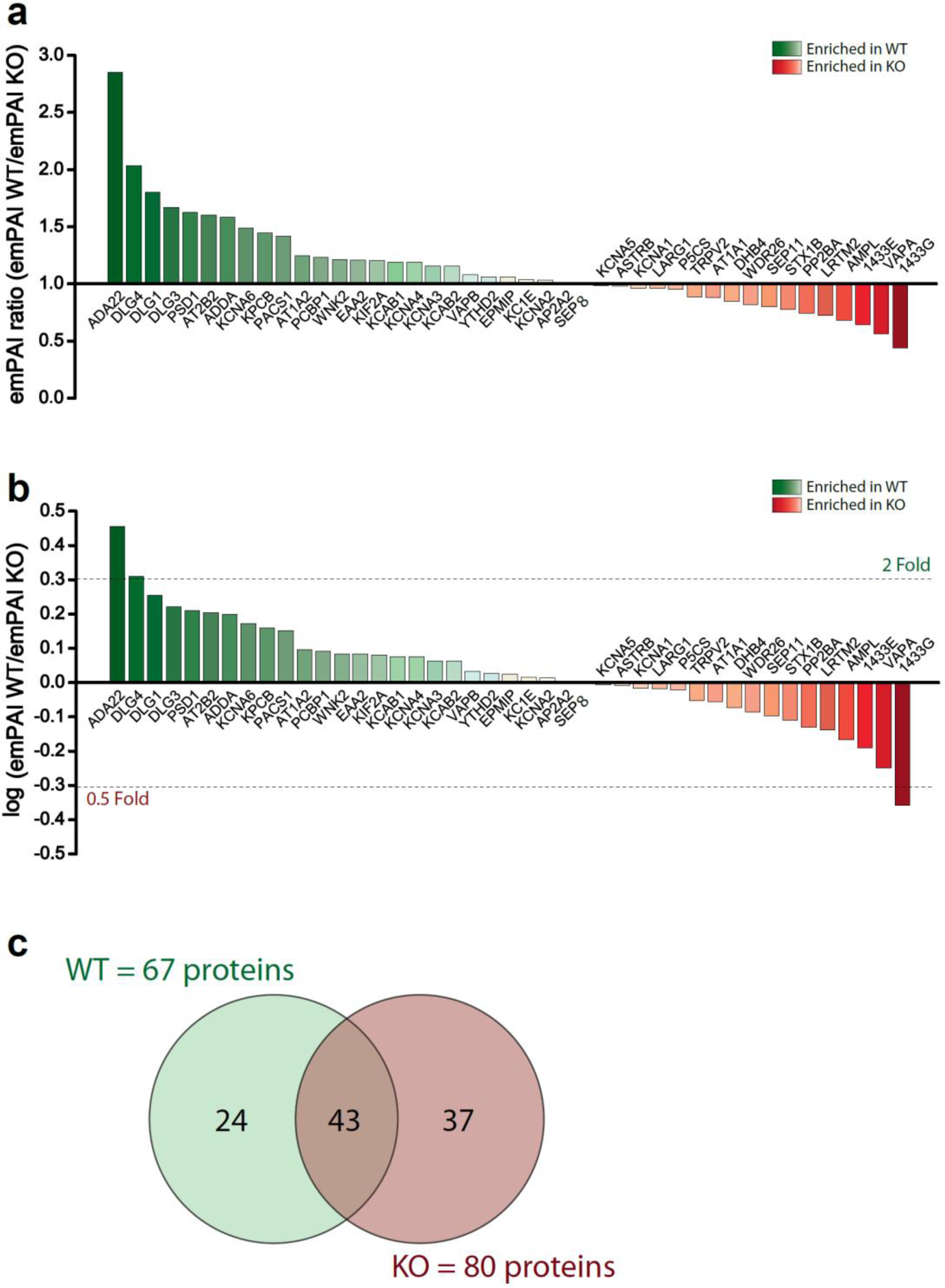
empAI ratios histogram of Kv_1_ partners in WT and *Lgi1^-/-^*. a) Principle up and downregulated Kv_1_ partners in WT and *Lgi1^-/-^*. b) Histograms of the log of ratios differences allowing to identify at a glance the fold change in partners enrichment. c) Venns diagram of Kv_1_ partners in this study.

**Figure S5.**
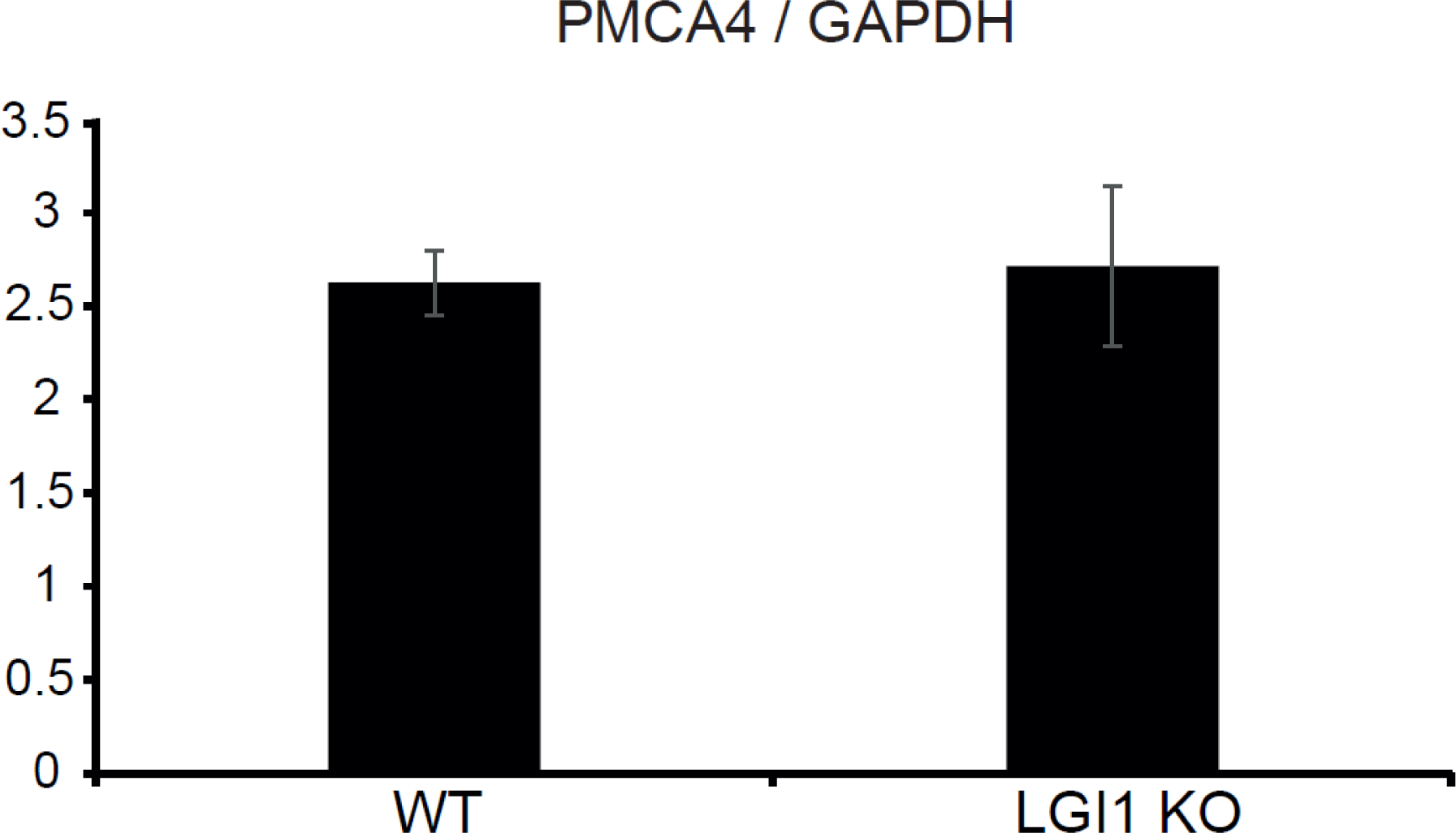
PMCA4/AT2B4 expression levels are not modified in *Lgi1^-/-^*. The expression level of PMCA4 was probed by Western blot on brain homogenates of WT and *Lgi1^-/-^*. Expression levels normalized to GAPDH signal are represented in histograms WT 2.62 ± 0.17 vs *Lgi1^-/-^* 2.71± 0.43 (p = 0.47152; n = 6; Two-tailed Mann-Whitney U test).

**Figure S6.**
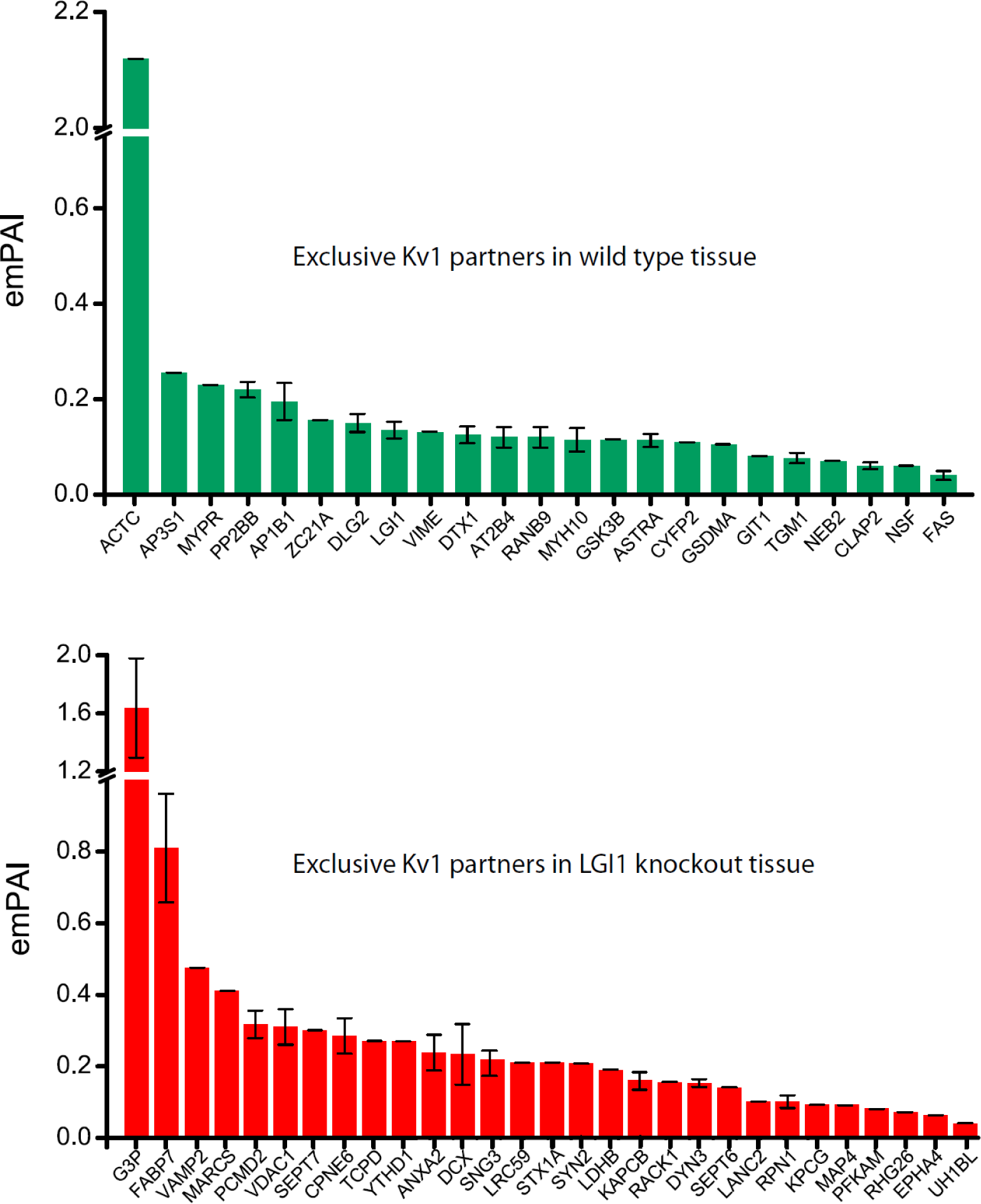
Exclusive Kv_1.2_ partners in WT and *Lgi1^-/-^.* Histograms showing exclusive immunoprecipitated protein partners of Kv_1.2_ from hippocampal homogenates in WT (top, green bars) and in *Lgi1*^-/-^ (bottom, red bars).

**Table S1.**
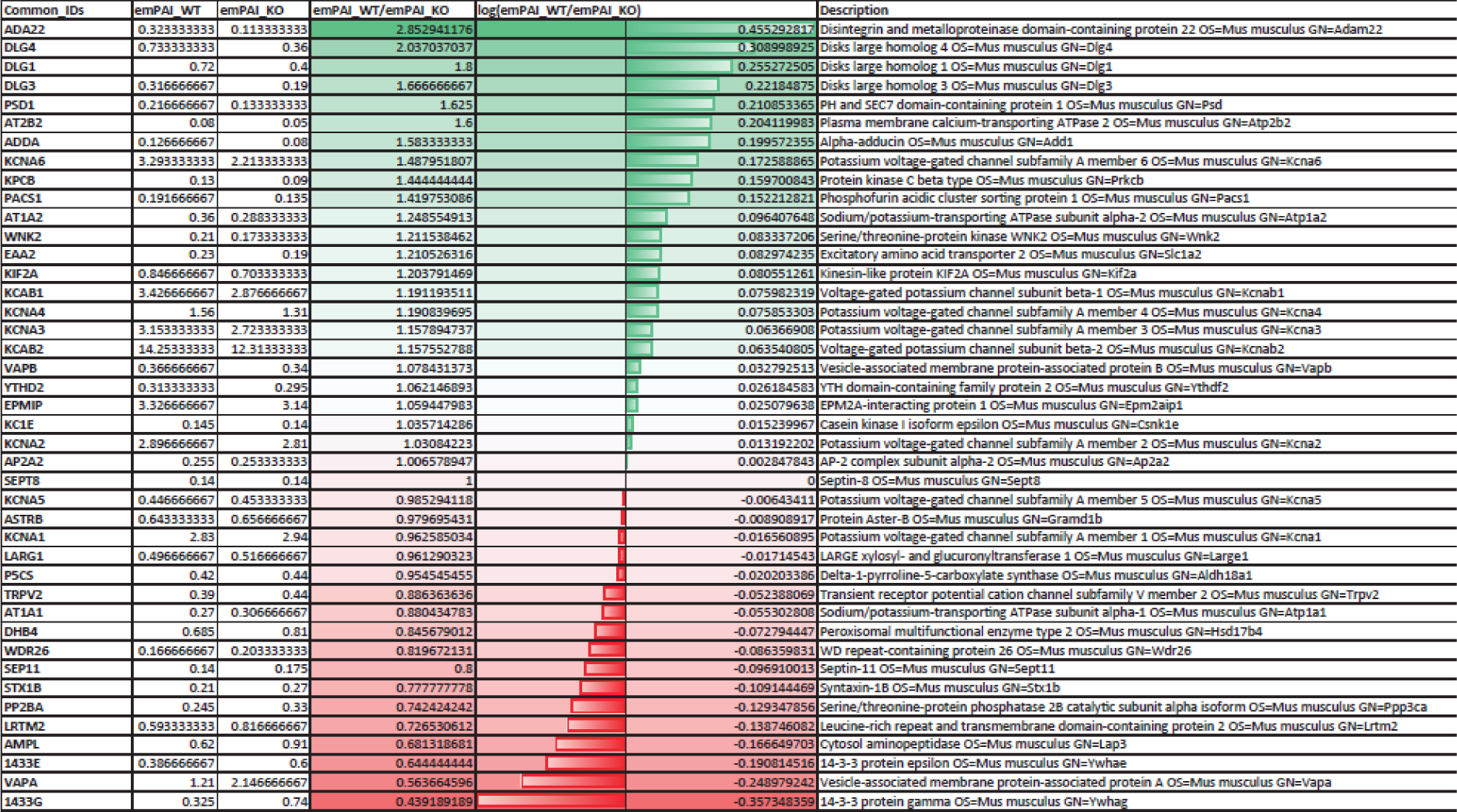
Common panners in WT and Lgi1 knock out background.

**Table S2.**
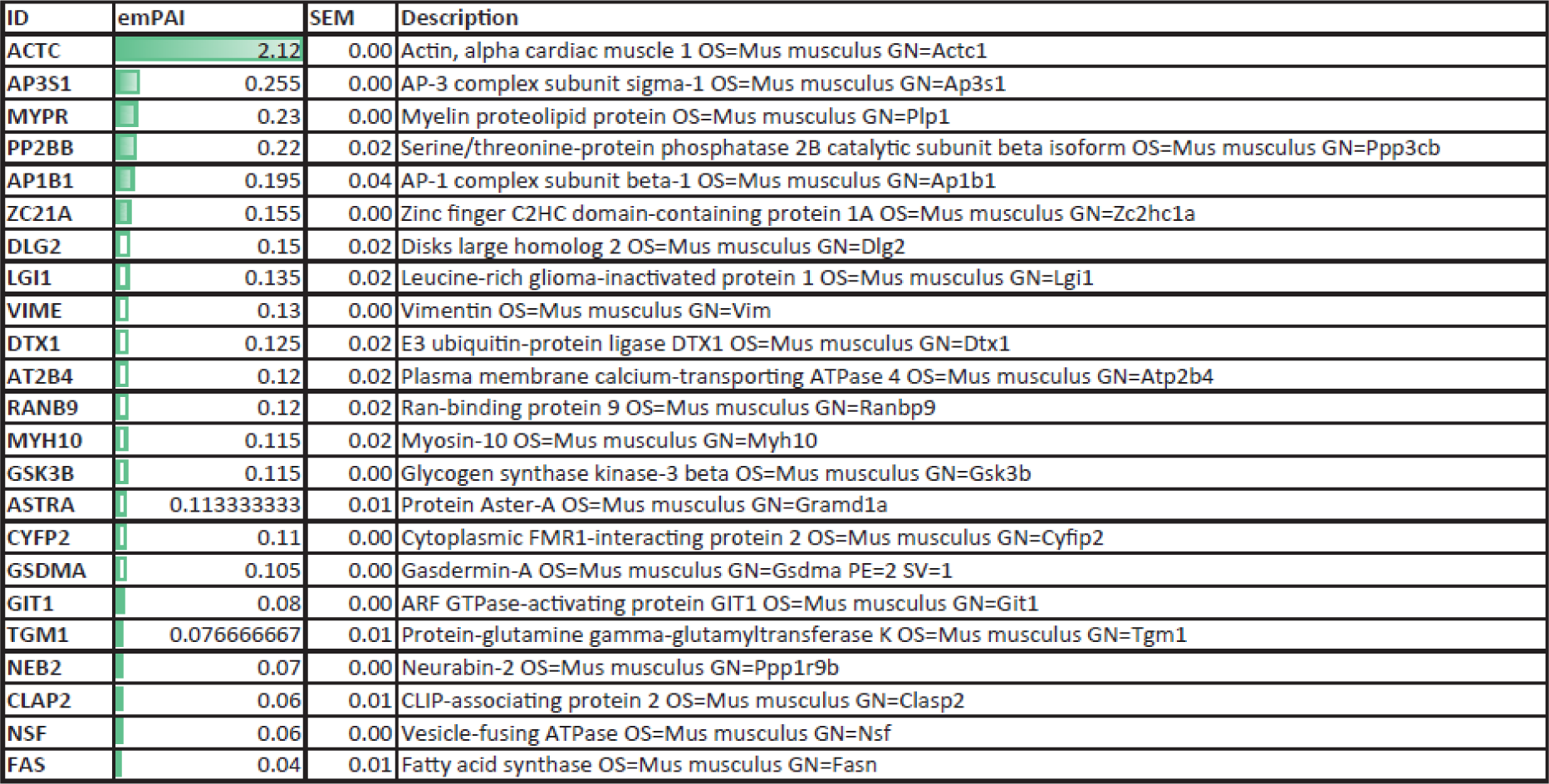
Specific Kv1.2 partners in WT background.

**Table S3.**
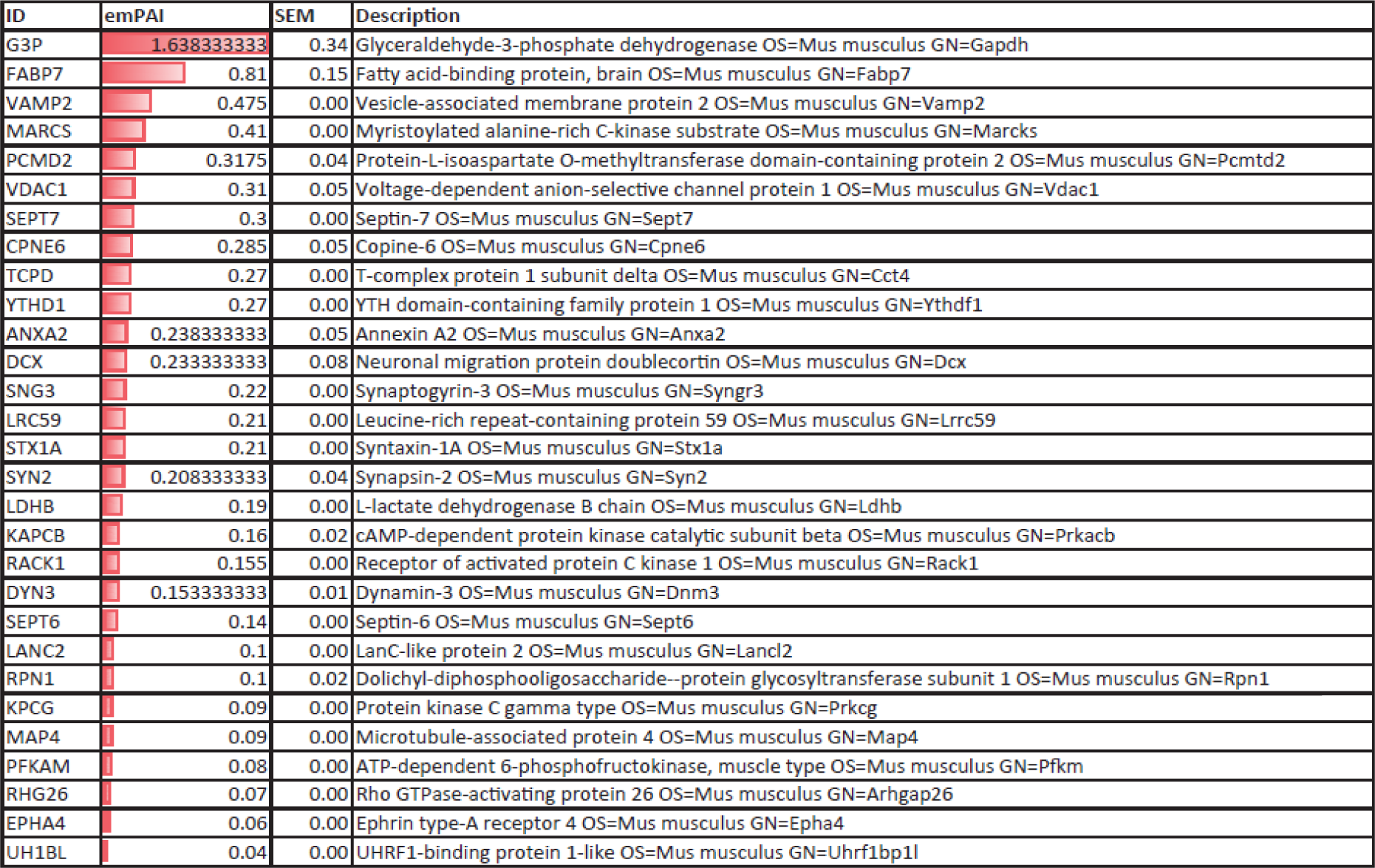
Specific Kv1.2 partners in Lgi1 know out background.

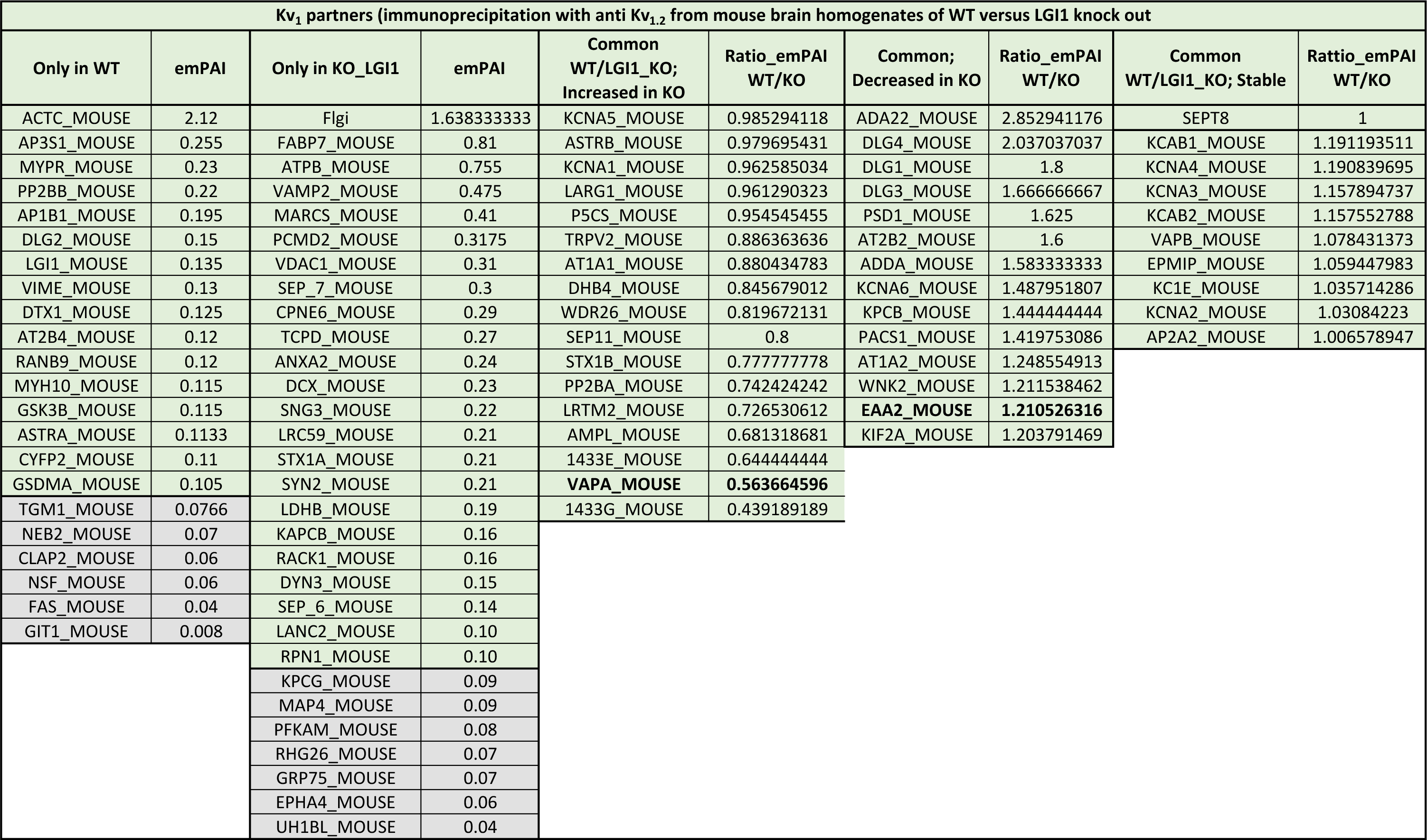
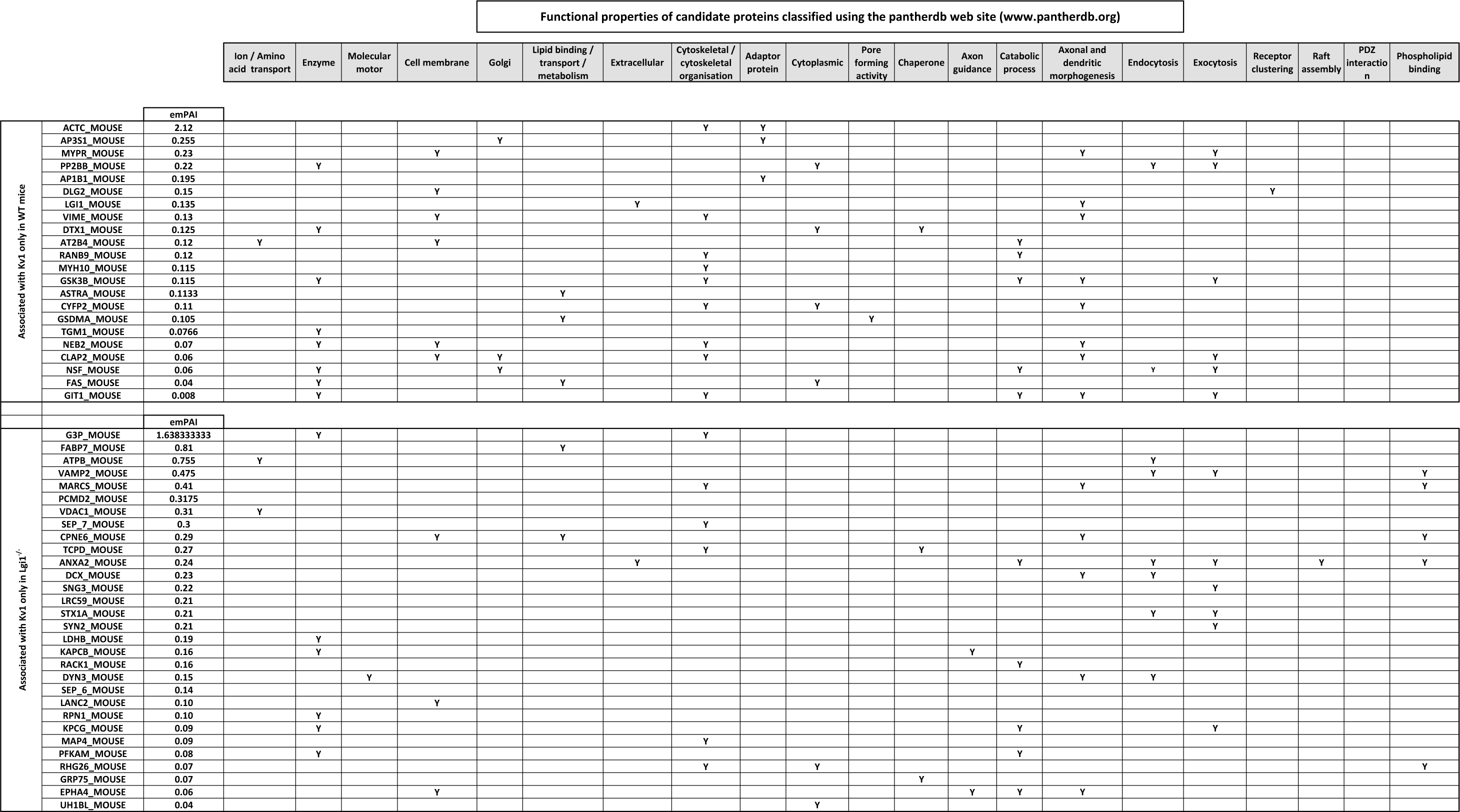
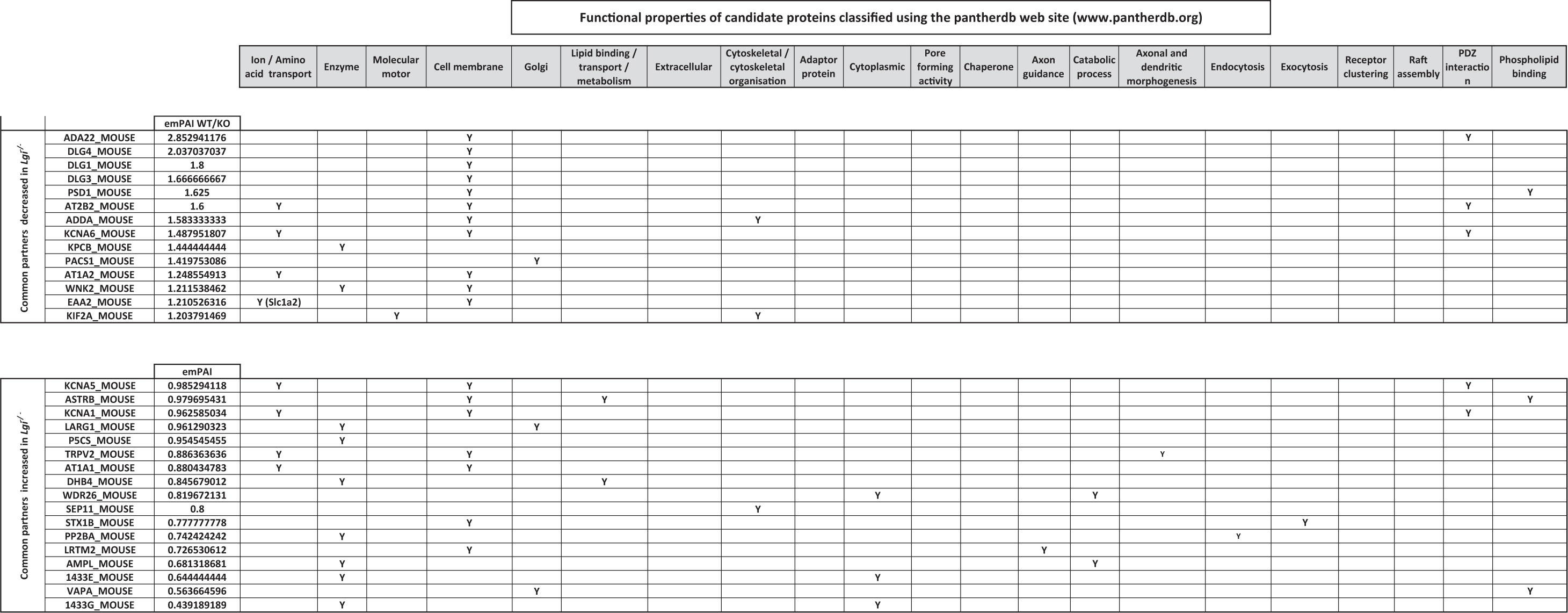
Supplementary sheet.

